# The wondrous and worrying diversity of the N‐glycans of *Chlorella* food supplements

**DOI:** 10.1101/2024.06.04.597335

**Authors:** Réka Mócsai, Johannes Helm, Karin Polacsek, Johannes Stadlmann, Friedrich Altmann

**Affiliations:** Department of Chemistry, BOKU University, Muthgasse 18, Vienna, Austria

**Keywords:** Microalgae, Chlorella, N‐glycan, glycoprotein, dietary supplements

## Abstract

N-glycans have recently emerged as highly varied elements of *Chlorella* strains and products. Four years and many samples later, the ever-growing N-glycan diversity shall be revisited in the light of concepts of species definition and product authenticity. N-glycans of commercial products were analyzed by matrix-assisted time-of-flight mass spectrometry (MALDI-TOF MS) supported by chromatography on porous graphitic carbon with mass spectrometric detection. While 36% of 172 products were labeled *C. vulgaris*, only few had matching N-glycan patterns. 5 and 20 % of the products matched with *C. sorokiniana* strains SAG 211-8k and SAG 211-34, respectively, which, however, carry entirely different structures. 41 % presented with four frequently occurring glyco-types while 26 % of the samples showed unique or rare N-glycan patterns. The rest presented what could be taken as a *C. vulgaris* type N-glycan pattern. Identical masses derive from different structures in many cases. By no means do we want to question the presumed health benefits of the products or the honest intentions of manufacturers. We rather wish to raise awareness for the fascinating but also worrying variety of microalgal N-glycans and suggest it as a means for defining product identity and taxonomic assignments.

## 1. Introduction

*Chlorella*, small green eukaryotic cells with hardly any obvious visual feature that aids discrimination of genera and species^1 2 3 4 5^. Even though, 42 genera within the family of *Chlorellaceae* are listed in the NCBI taxonomy repository and the genus *Chlorella* is split up into 25 species. The difficulty of unequivocal classification of microalgae is demonstrated by rather frequent re-classification events that sometimes even jump across the class borders as in the case of the late *Chlorella fusca*, which is now found as the former name of four different species (http://sagdb.uni-goettingen.de) from the class of *Chlorophyceae* ^6^. For the newcomer: this is exactly the class that does not comprise the *Chlorellaceae* family with its best-known character *Chlorella*.

These uncertainties did not prevent scientists from evolving an awe-inspiring multitude of applications of *Chlorella* and other microalgae. In 2022, 671 PubMed entries contained the term *Chlorella* in a highly diverse array of contexts reviewed *e*.*g*. by Abreu et al. ^7^. Frequent items are the production of biomass and lipids, sometimes with the scope of biodiesel production. These low-value applications are often connected to objectives such as waste water treatment, heavy metal, nitrogen or phosphorus depletion and – last but not least – carbon sequestration. A remarkable niche application is the expression of antigenic protein for the vaccination of shrimps ^8^. Higher priced products for human consumption comprise dyes such as astaxanthin or phycocyanin, unsaturated fatty acids (omega-3 fatty acids), vegan protein and vitamins ^9^. A natural, vegan source of vitamin B_12_ certainly is of interest, whereby one should be cautious as older determination methods could not discriminate true vitamin B_12_ from its biologically inactive pseudo-vitamin that prevails in *Spirulina* ^10^. Rather varied contents of B_12_ were recently found in a panel of *Chlorella* samples ^10^ and this may well reflect the varied nature of the microalgae constituting the products as visualized by their differing N-glycosylation. Even though, the cobalamin content appears to be the best substantiated one of the many health claims for *Chlorella* dietary supplements. In contrast, a scientific basis for the likewise often acclaimed benefit of the high chlorophyll content of microalgae has not been brought to the attention of the authors of the present treatise.

Recently, we found that different culture collection lines as well as *Chlorella* nutraceuticals contained a range of different N-glycans and N-glycan patterns when viewed by mass spectrometry ^11^. Very few products exhibited a *C. vulgaris* pattern with methylated oligomannosidic N-glycans only, several products displayed a *C. sorokiniana* ^12^ and others a “FACHB31” pattern ^13^ with very characteristic major glycans (Figure 1). Apart from that, the 80 products split up into at least 10 different glyco-types ^11^. Notably, 22 of these products were labeled as *C. pyrenoidosa*, which is an invalid species designation for years already ^6, 14^. Even though, PubMed contains 36 publications on *C. pyrenoidosa* as of April 2, 2024, many of which without an explicit strain number.

**Figure 1.**
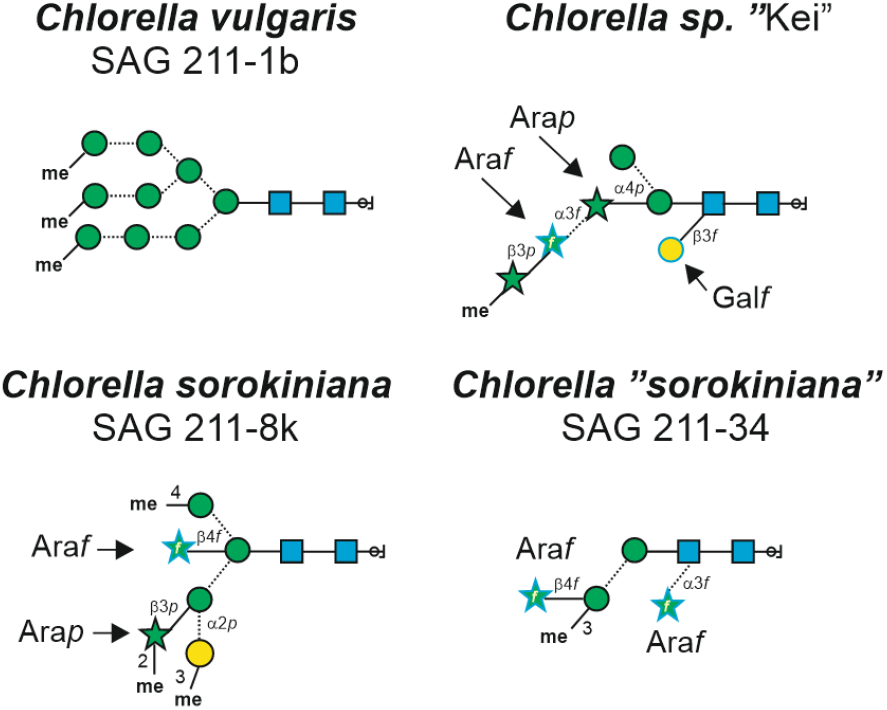
Structures of N-glycans from *Chlorella* (clade) strains known to date ^12, 13, 15^.

The development of this field deserves interest and therefore we ordered and analyzed another 90 *Chlorella* products, several of them re-purchases. A small number of frequently found glycan patterns re-emerged together with a remarkable number of novel patterns. We try to bring some order into the many glycan patterns as surfaced by matrix-assisted laser-desorption time-of-flight mass spectrometry (MALDI-TOF MS) together with a procedure discerning isobaric N-glycans with the help of retention-time standardized liquid chromatography electrospray-ionization mass spectrometry on porous graphitic carbon (PGC-LC-MS). While the results of this study raise doubts about the current practice of *Chlorella* strains designation, they also point at strategies for improved product characterization and control.

## 2. Materials and Methods

### 2.1 Chlorella samples

Commercial *Chlorella* food supplements were ordered on-line or bought at local stores. A list of product names, principal distributors and lot numbers is found as Table S1. Live cultures of culture collection strains were grown photoautotrophically in Bold’s basal medium (without antibiotics) as described ^15^.

### 2.2 N-glycan preparation and analysis by MALDI-TOF MS and MS/MS

N-glycans were prepared by the differential application of cation exchange before and after digestion with peptide:N-glycosidase A (PNGase A; Europa Bioproducts, www.europa-bioproducts.com) as described ^11, 15, 16^. Free N-glycans were co-crystallized with 2,5-dihydroxybenzoic acid and analyzed with an Autoflex MALDI-TOF MS (Bruker, www.bruker.com) in positive reflectron mode. All spectra were re-calibrated using oligomannosidic glycans as internal standards. Almost all signals qualified as representing N-glycans according to their composition of hexose, HexNAc, pentose and methyl (which together might mask a deoxyhexose) residues as as calculated by the spread sheet “algae mass converter” (Table S2). A second criterion was the embedding in a cluster of similar compositions differing e.g., in the number of methyl groups. Fragment spectra were obtained by laser induced fragmentation (LIF) in “LIFT” mode and tentatively interpreted using the spread sheet Excel tool (Table S3).

### 2.3 Analysis by porous graphitic carbon liquid chromatography (PGC-LC-MS)

For analytical purposes, the glycans were reduced with NaBH_4_, desalted by passage over graphitized carbon cartridges (ThermoFisher Scientific, www.thermofisher.com) and subjected to LC-MS with a PGC column (0.32 mm; 150 mm) connected to an amazon ion trap (Bruker) operating in data-dependent acquisition mode ^11^. The PGC column was eluted with 65 mM ammonium formate at pH 3.0 and a gradient from 8 to 60% acetonitrile in 50 min at a flow rate of 6 μl min^-1^ at 30°C. Retention times were normalized with the help of internal standards as described ^17^. However, adapted to the current task, unlabeled porcine brain N-glycans and ^13^C_4_-A^4^A^4^F^6^, which was prepared from porcine fibrin exploiting the recently discovered de-*N*-acetylating potency of hydrazine hydrate ^18^, were used as internal standards. The respective measured retention times were converted to virtual minutes (v-min).

## 3. RESULTS

### 3.1. A roadmap through the glycan maze

During the last four years our collection of commercial *Chlorella* samples has been extended to now comprise 172 different samples, all with different brand names in vividly colored, attractive packaging. The samples came from vendors mostly from Europe and mostly from resellers, who largely lacked inclination to reveal the actual producer of a given lot. Thus, unfortunately the source of the products remains in the dark. Products with very similar patterns may or may not stem from the same production site. On the other hand, it may happen – deliberately or not – that the production strain in a particular site changes over time. This may be the underlying reason for the observation that different lots from the same vendor exhibited different glycan patterns (13 out of 19 re-ordered samples gave a clearly different pattern; this data is, however, not explicitly revealed herein). The same situation could however result from a change of supplier. A further source of confusion could be mixed production strains or blending of previously “pure” products. Despite these regrettable limitations of the current study, the matter deserves attention particularly in the light of food identity and authenticity.

Table S1 lists the products, their sources, lot numbers and the stated *Chlorella* species. The following chapters try to structure the maze of variations and peculiarities. At first, we will deal with glycan patterns matching those of established culture collection strains. The remaining 76 % of samples and their patterns will be split into four larger, several smaller groups and finally in apparently unique patterns. This grouping is essentially based on mass spectrometry of N-glycans with some assistance from (18S)-ITS1-5.8S-ITS2-(23S) rRNA barcoding. In the following, relevant characteristics of the glycan pattern groups will be presented.

### 3.2 Vendors’ designations of *Chlorella* products

The food market in Europe is regulated. The microalgae species that can be freely marketed as they do not fall under the Novel Food Regulation 2283/2015 of the European Union are *C. vulgaris, C. pyrenoidosa* and *C. luteoviridis* (now renamed as *Jaagichlorella luteoviridis*) (https://www.algae-novel-food.com/output/algae-novel-food/download.pdf). Among our collection of 172 samples, almost half of the products did not provide a species name, while the other half split up rather equally into *C. vulgaris* and *C. pyrenoidosa* (Table S1). As our internal specimen numbers were assigned chronologically over a period of eight years, the use of species terms over time can be scheduled (Figure 2). A slight decrease of the popularity of the term *C. pyrenoidosa* can be sensed, which may be attributed to ever more producers or retailers realizing that this species name was officially abandoned by taxonomists and type strain collections ^6^. One product was designated *C. sorokiniana*. This product came from the United States where other regulations apply than in Europe.

**Figure 2.**
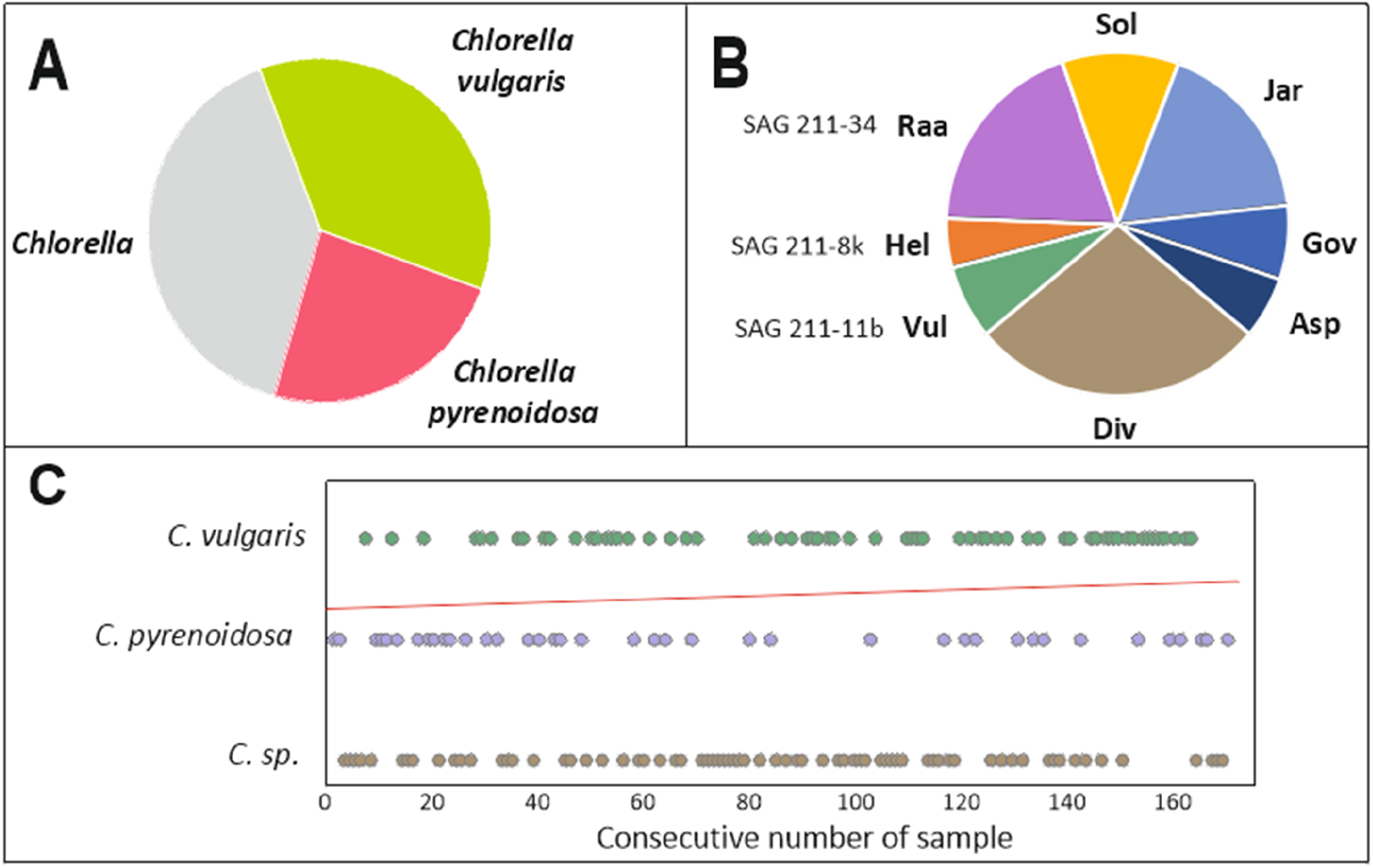
Panel **A** illustrates the distribution of product designations. Panel **B** depicts the frequencies of the glyco-patterns found in the study group of 172 *Chlorella* products. The three letter abbreviations were introduced recently ^11^ and are explained in the following chapters and in Table 1. “Div” comprises all unique or rare patterns possibly including a few “misdiagnosed” ones. “Vul” bunches together spectra that at least remotely resemble the previously described *C. vulgaris* glycosylation pattern. Panel **C** shows the distribution of the product designations over sample numbers, which roughly reflect purchase time during the last eight years. The red line depicts the trend of the frequency of the two species designations.

### 3.3 Structuring the pandemonium of MALDI-TOF MS N-glycan patterns

In this study, we focus on the N-glycan spectra derived from the various products. Due to the extraction and detection procedures, these spectra display neutral N-glycans only. However, hitherto, no acidic or zwitterionic N-glycans have been found in algae. The combination of varying numbers of hexoses, pentoses, methyl-groups and even *N*-acetylhexosamines (HexNAc) leads to a vast variety of mass values. Note, that this study does not identify deoxyhexoses, which are isobaric to the combination of a pentose and a methyl-group and which have so far not been found in Chlorella clade algae ^11^. From the 132 counted mass values, 36 represent oligomannosidic glycans with and without methyl-groups, 91 comprise pentose-containing N-glycans and 18 even contain three to four HexNAcs. The exact masses and examples of how samples could be depicted in a QR code like manner are shown in (Table S4) and exemplary in Figure 3. Note, that in the following text and in main text figures, often only nominal masses are given as the addition of 1, 2 or more digits behind the comma would be as impressive as pointless. Compositions can be derived from masses by the spread sheet “algae mass converter” (Table S2).

**Table 1.**
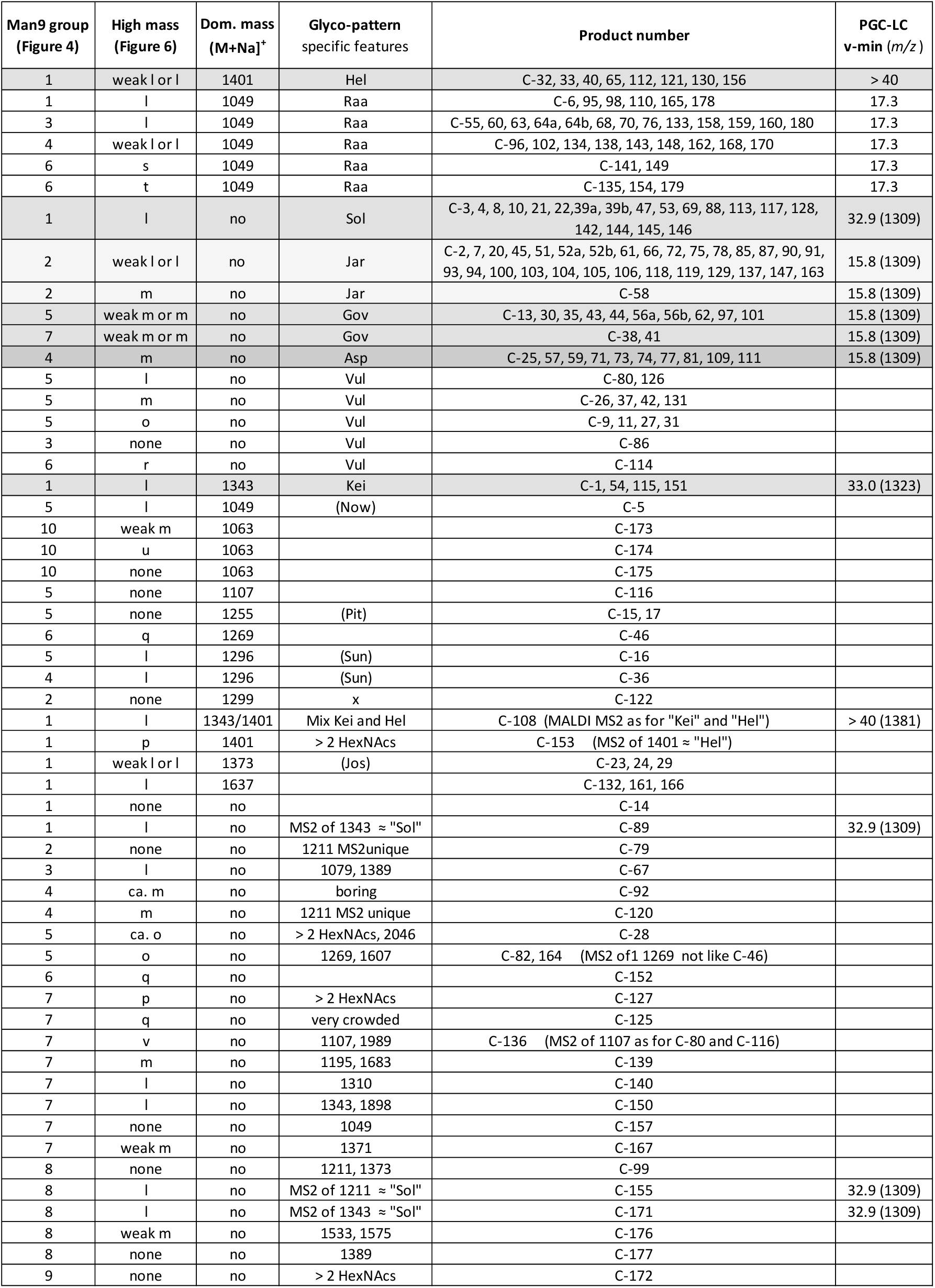
Glyco-patterns of *Chlorella* products. N-glycan patterns generated by MALDI-TOF MS were grouped according to various features to the best of the authors abilities. Example spectra of glyco-patterns are found in Figure S2, all other spectra in Figure S6. Glyco-patterns put in brackets were mentioned previously ^11^ but did not re-occur in new samples.

| Man9 group<br>(Figure 4) | High mass<br>(Figure 6) | Dom. mass<br>(M+Na)<sup>+</sup> | Glyco-pattern<br>specific features | Product number | PGC-LC<br>v-min (m/z) |
| --- | --- | --- | --- | --- | --- |
| 1 | weak l or l | 1401 | Hel | C-32, 33, 40, 65, 112, 121, 130, 156 | > 40 |
| 1 | l | 1049 | Raa | C-6, 95, 98, 110, 165, 178 | 17.3 |
| 3 | l | 1049 | Raa | C-55, 60, 63, 64a, 64b, 68, 70, 76, 133, 158, 159, 160, 180 | 17.3 |
| 4 | weak l or l | 1049 | Raa | C-96, 102, 134, 138, 143, 148, 162, 168, 170 | 17.3 |
| 6 | s | 1049 | Raa | C-141, 149 | 17.3 |
| 6 | t | 1049 | Raa | C-135, 154, 179 | 17.3 |
| 1 | l | no | Sol | C-3, 4, 8, 10, 21, 22, 39a, 39b, 47, 53, 69, 88, 113, 117, 128, 142, 144, 145, 146 | 32.9 (1309) |
| 2 | weak l or l | no | Jar | C-2, 7, 20, 45, 51, 52a, 52b, 61, 66, 72, 75, 78, 85, 87, 90, 91, 93, 94, 100, 103, 104, 105, 106, 118, 119, 129, 137, 147, 163 | 15.8 (1309) |
| 2 | m | no | Jar | C-58 | 15.8 (1309) |
| 5 | weak m or m | no | Gov | C-13, 30, 35, 43, 44, 56a, 56b, 62, 97, 101 | 15.8 (1309) |
| 7 | weak m or m | no | Gov | C-38, 41 | 15.8 (1309) |
| 4 | m | no | Asp | C-25, 57, 59, 71, 73, 74, 77, 81, 109, 111 | 15.8 (1309) |
| 5 | l | no | Vul | C-80, 126 |  |
| 5 | m | no | Vul | C-26, 37, 42, 131 |  |
| 5 | o | no | Vul | C-9, 11, 27, 31 |  |
| 3 | none | no | Vul | C-86 |  |
| 6 | r | no | Vul | C-114 |  |
| 1 | l | 1343 | Kei | C-1, 54, 115, 151 | 33.0 (1323) |
| 5 | l | 1049 | (Now) | C-5 |  |
| 10 | weak m | 1063 |  | C-173 |  |
| 10 | u | 1063 |  | C-174 |  |
| 10 | none | 1063 |  | C-175 |  |
| 5 | none | 1107 |  | C-116 |  |
| 5 | none | 1255 | (Pit) | C-15, 17 |  |
| 6 | q | 1269 |  | C-46 |  |
| 5 | l | 1296 | (Sun) | C-16 |  |
| 4 | l | 1296 | (Sun) | C-36 |  |
| 2 | none | 1299 | x | C-122 |  |
| 1 | l | 1343/1401 | Mix Kei and Hel | C-108 (MALDI MS2 as for "Kei" and "Hel") | > 40 (1381) |
| 1 | p | 1401 | > 2 HexNAcs | C-153 (MS2 of 1401 ≈ "Hel") |  |
| 1 | weak l or l | 1373 | (Jos) | C-23, 24, 29 |  |
| 1 | l | 1637 |  | C-132, 161, 166 |  |
| 1 | none | no |  | C-14 |  |
| 1 | l | no | MS2 of 1343 ≈ "Sol" | C-89 | 32.9 (1309) |
| 2 | none | no | 1211 MS2unique | C-79 |  |
| 3 | l | no | 1079, 1389 | C-67 |  |
| 4 | ca. m | no | boring | C-92 |  |
| 4 | m | no | 1211 MS2 unique | C-120 |  |
| 5 | ca. o | no | > 2 HexNAcs, 2046 | C-28 |  |
| 5 | o | no | 1269, 1607 | C-82, 164 (MS2 of 1269 not like C-46) |  |
| 6 | q | no |  | C-152 |  |
| 7 | p | no | > 2 HexNAcs | C-127 |  |
| 7 | q | no | very crowded | C-125 |  |
| 7 | v | no | 1107, 1989 | C-136 (MS2 of 1107 as for C-80 and C-116) |  |
| 7 | m | no | 1195, 1683 | C-139 |  |
| 7 | l | no | 1310 | C-140 |  |
| 7 | l | no | 1343, 1898 | C-150 |  |
| 7 | none | no | 1049 | C-157 |  |
| 7 | weak m | no | 1371 | C-167 |  |
| 8 | none | no | 1211, 1373 | C-99 |  |
| 8 | l | no | MS2 of 1211 ≈ "Sol" | C-155 | 32.9 (1309) |
| 8 | l | no | MS2 of 1343 ≈ "Sol" | C-171 | 32.9 (1309) |
| 8 | weak m | no | 1533, 1575 | C-176 |  |
| 8 | none | no | 1389 | C-177 |  |
| 9 | none | no | > 2 HexNAcs | C-172 |  |

**Figure 3.**
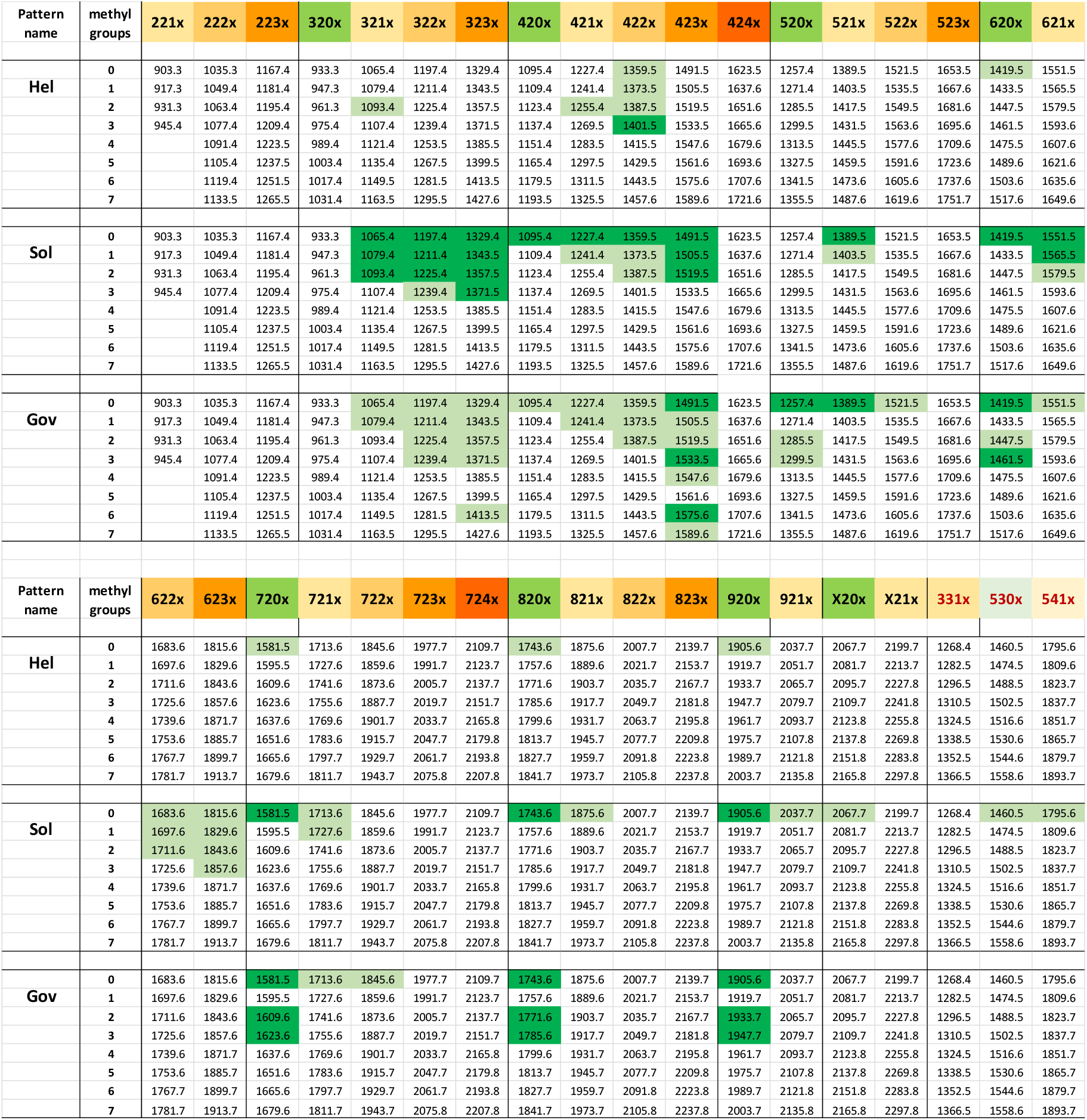
Microalga mass map with three arbitrarily selected examples. The colored upper rows give the number of hexoses, HexNAcs and pentose residues in a structure. The rows below list the *m/z* values of [M+Na]^+^ ions with increasing numbers of methyl groups. Light and dark green cell colors represent smaller (approximately 5 to 30 % of maximal peak height) and larger peaks. A more comprehensive and depiction is available as Table S4.

Depiction of full range spectra would necessarily conceal details. Therefore, just a few sections of particular diversity are shown. One shall be the Man9 region showing ten characteristic methylation patterns (Figure 4). A second example shall be the region *m/z =* 1300-1600 (Figure 5). As a third and last example, the high-mass region was chosen, which – beyond being impressively divers – confirms the ability of microalgae to generate glycans with up to four HexNAc residues (Figure 6).

**Figure 4.**
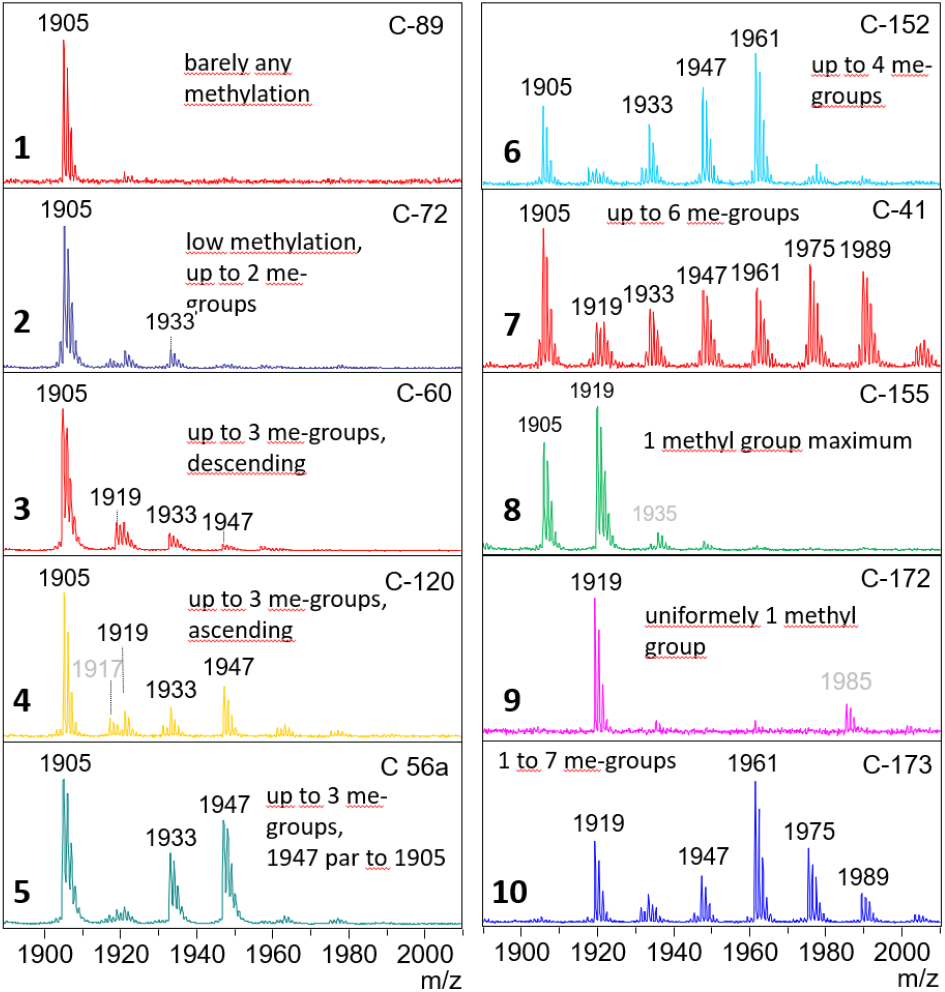
Ten characteristic patterns of O-methylation of the oligomannosidic N-glycan Man9 (Man_9_GlcNAc_2_). The numbers in the upper right corners refer to the respective *Chlorella* product (Table S1). Peaks are labeled with nominal *m/z* of the [M+Na]^+^ ions. Note, that in this and all following spectra large peaks can be followed by a [M+K]^+^ satellite.

**Figure 5.**
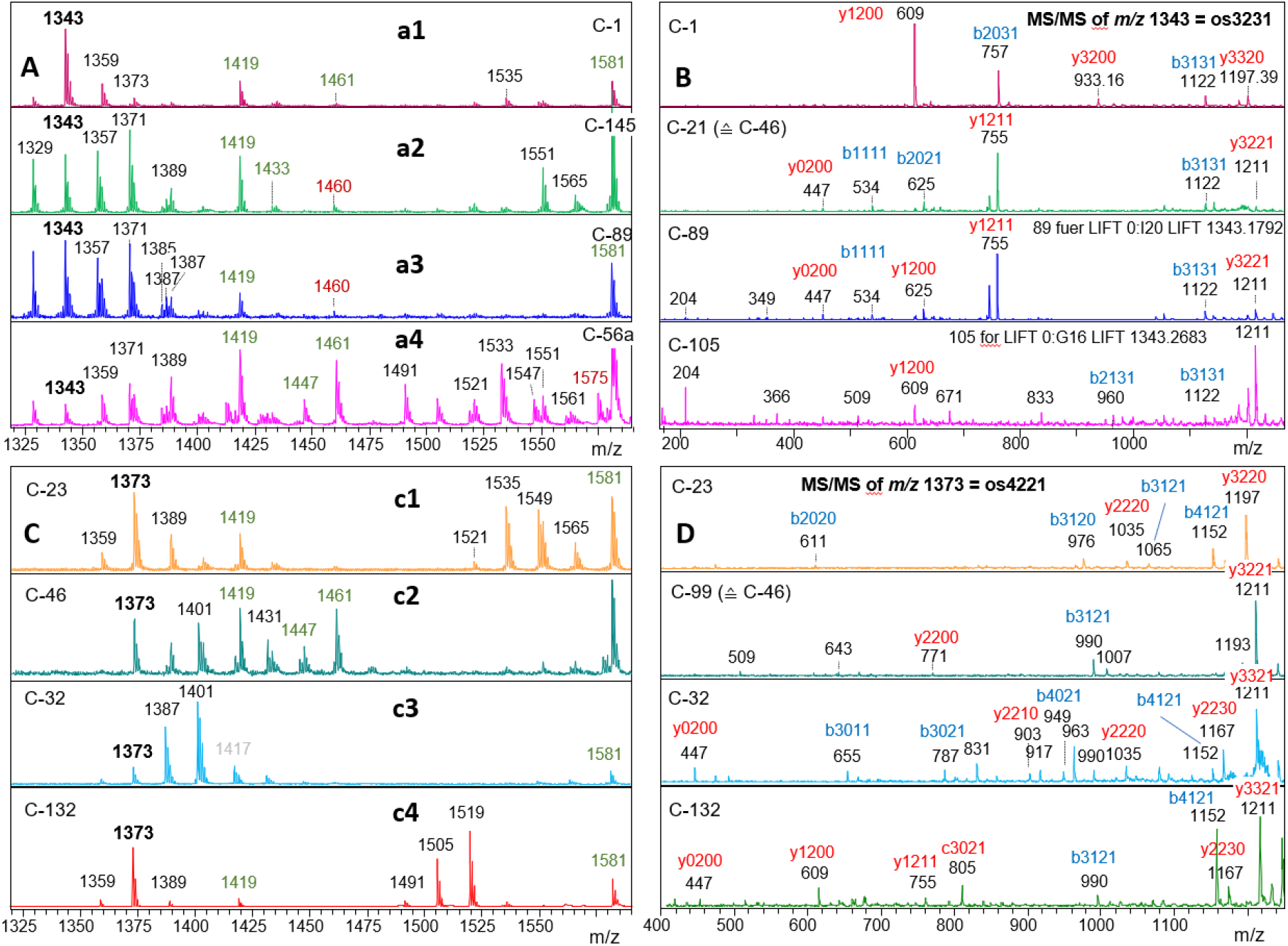
Seven distinct patterns of the medium mass range of microalgal N-glycans. MS1 spectra are shown on the left, LIF-MS/MS spectra of *m/z* = 1343.4 and 1373.4 on the right side. Fragments are tentatively assigned as y (in red) and b (and c; in blue) ions with the numbers of hexose, HexNAc, pentose and methyl constituents. The MS/MS spectra indicate three different structures both for *m/z* = 1343.4 and 1373.4.

**Figure 6.**
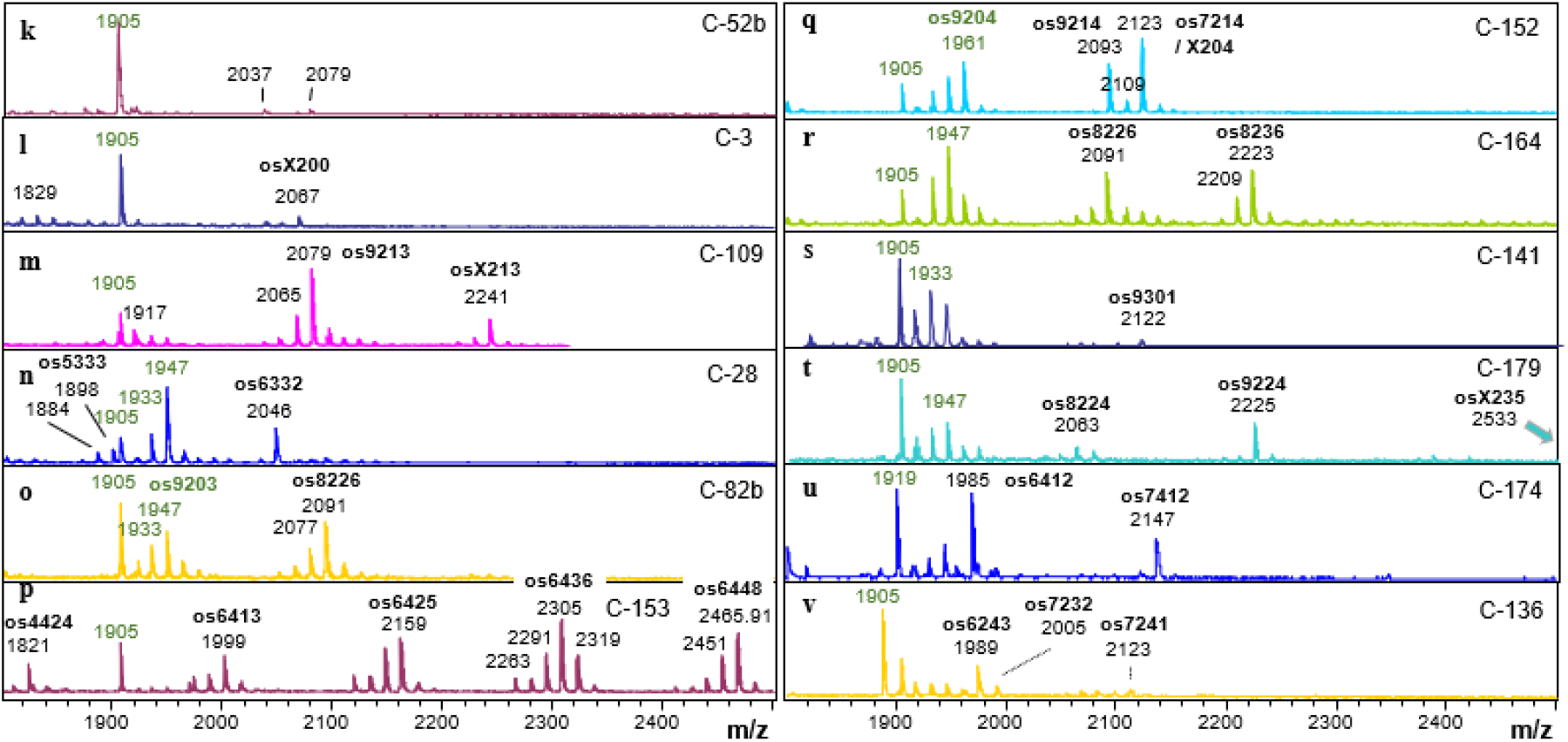
Twelve peculiar spectra of the high mass region of *Chlorella* N-glycans. Oligosaccharides (os) are characterized by the numbers of hexose, HexNAc, pentose and methyl constituents, whereby X stands for 10.

A fourth and highly helpful criterium is the occurrence of a dominant “lead” mass (primarily *m/z* 1049, 1343, and 1401) that can be used to cluster product spectra. By a happy coincidence, the *m/z* values 1049 and 1401 represent the outstanding glycans of products for which culture collection type strains are known ^12, 13^.

### 3.4 Commercial samples with a *C. vulgaris* pattern

With more than one third of all products being designated as *C. vulgaris*, one could expect an equally large number of samples to present an N-glycan pattern similar to that of the strains SAG211-11b, SAG 211-8l, UTEX235 ^15^ plus now SAG 211-80 and SAG 211-8m (Figure S1). The hallmark of these *C. vulgaris* strains is the exclusive presence of oligomannosidic N-glycans of which the larger part is equipped with up to three O-methyl groups ^15^. This “type V” pattern of Man9 glycans was found in 14 % of all samples. However, the majority of these products exhibited considerable amounts of additional peaks containing pentoses, possibly also deoxyhexoses and again methyl groups. Only 12 products could be accepted as having a rather vulgaris-like appearance. For one sample (C-126) DNA barcoding was performed and it corroborated the classification as *C. vulgaris* (Supplementary sequences). However, almost exclusively, did this samples contain additional smaller peaks representing “pentose-glycans” (Figure S2). We cannot define whether these extra peaks are a previously overlooked feature of *C. vulgaris* or result from minor contamination of the cultures with other algae strains. At any rate were these extra peaks highly varied. Another notable detail is that only two out of these 12 products carried the label *C. vulgaris*. Viewed from a different angle, 60 samples are labeled *C. vulgaris* but present a clearly different N-glycan pattern.

### 3.6 Samples with *C. sorokiniana* patterns

Three *C. sorokinian*a strains (the type strain SAG 211-8 k and also SAG 211-32 and UTEX 1230) carried one very dominant glycan with structural elements not seen before ^12^. To complicate the situation, two other “*C. sorokiniana*” strains (SAG-211-31, SAG211-34; and also FACHB-31 surprised with what almost resembled a number twister of the major peak’s mass (Figure S1). Instead to of *m/z* = 1401, the dominant nominal mass here was 1049. At first sight, the small glycans could be a precursors or a degradation product of the larger one, but the structures of both N-glycans do not present any resemblance ^13^. One can infer the number of specialized, unusual glycosyltransferases required to generate the one or the other glycan. With 6 and 4 such transferases needed to be evolved by evolution just for these “*C. sorokiniana*” variants, we can no longer assume that these two glyco-types fit under the roof of the same species name. As for the moment, we have to stick to these names, we will refer to these glyco-variants as “Hel” (*C. sorokiniana* with MALDI *m/z* = 1401) and “Raa” (*C. sorokiniana* and FACHB-31 with MALDI *m/z* = 1049) patterns as in the previous publication ^11^. The particular spice comes from the fact that while only one product was labeled *C. sorokiniana*; eight had a “Hel” pattern and no less than 34 had a “Raa” glycosylation (Figure 2, Table 1). Ironically, the one product labeled *C. sorokiniana* did not fit into either group. A further distinction of the “Raa” (*m/z* = 1049) group can be made based upon the methylation of Man9 (Table 1). Whatever the reason for these differences might be, LIF-MS/MS of the *m/z* = 1049 peak in all of these samples indicated identical glycan structure (Figure S3). Caution must, however, be exerted as the solitary sample C-5 gave a rather similar MS/MS spectrum despite a totally different structure ^13^. LC--MS of selected samples advocated the notion of identical glycan structure (see separate chapter on LC-MS below) (Figure S4).

### 3.7 Frequently found “orphan” glyco-types

The term “orphan” encompasses glyco-types for which no culture collection strain has so far been identified. In the bewildering diversity of the often very crowded MALDI-TOF MS spectra of *Chlorella* N-glycans, three groups of densely nested signals were found in a large number of samples. These groups started with the masses 1197, 1329 and 1491^1^ (Figure 5). These figures are the simplified, integer *m/z* values of the [M+Na]^+^ ions observed by MALDI-TOF MS of released and purified N-glycans. A daunting complexity results from the addition of one or more methyl groups (14 mass units each). In some patterns, the oligomannosidic N-glycans were likewise methylated and contributed to signal plurality (Figure 4). As if not complex enough, potassium ions cause 16 mass unit increments – usually, but not always, with peaks much smaller than the main [M+Na]^+^ peak.

To get firm ground under our feet, we at first figured out the possible composition of the mentioned masses. Our initial work had shown the hexoses mannose and galactose and the pentoses arabinose and xylose and, of course, HexNAc as the building blocks of the “club 1329” members ^11^. A glycan mass can then be annotated in terms of the number of hexoses, HexNAc and pentoses and the number of methyl groups. A MALDI reading can be easily converted to a composition using the “algae mass converter” (Table S2). The [M+Na]^+^ value 1329 as an example figures for a glycan with 3 hexoses, 2 HexNAcs (*i*.*e*. GlcNAc) and 3 pentoses and is annotated as os3230. *M/z* = 1343 then translates into os3231 and so on. The many combinations – realized or not by nature – can be listed in a spread sheet as shown in Figure 3 and Table S4.

At least 74 samples adhered to the “1197, 1329 and 1491 plus methyl-groups” schemes **a2** or **a3** shown in (Figure 5). The samples differed in the occurrence and type of OM methylation and masses beyond *m/z* 2000. The distinction of samples with no or very little OM methylation (Man9 type I and II in (Figure 4) could have been seen as futile were it not accompanied by the occurrence of *m/z* = 1460 (os5300) and 1795 (os5410) in the “Sol” group. Os5300 was shown to be the Man5GlcNAc that forms the pivotal transition point from oligomannosidic to complex-type N-glycans in land plants and animals ^11^. LIF-MS/MS of the *m/z* = 1343 peaks of samples exhibiting the **a1** to **a3** patterns substantiated the difference between patterns **a2** (“Sol”) and **a3** (“Jar”), as well as pattern **a1**, which is only found in the rare glyco-group “Kei” ^11, 12^. The **a3** glyco-pattern was also seen in the more complex glyco-groups “Gov” and “Asp”, which unfortunately defied LIF-MS/MS. Their close relation to “Jar” was, however, substantiated by LC-ESI-MS (see separate chapter below).

The small “Kei” group with its predominant *m/z* = 1343 shall be added here mainly because it’s major glycan structure has been elucidated ^12^.

At this stage we had categorized 76 % of the 172 products as either “Hel” (*m/z* = 1401), “Raa” (*m/z* = 1049), “Kei” (*m/z* = 1343), or the more complex glyco-groups “Sol”, or “Jar-Gov-Asp” (Figure 1). The rest, unfortunately, splits up into a rather intricate variety of glycan-patterns.

### 3.8 Defying classification

24 % of the products could not – or at least not at first and second sight – be associated with any of the glyco-groups described above. Their features are listed in Table 1, whereby the primary sorting criterium was the occurrence of a dominant MALDI-TOF MS peak. Eleven different *m/z* values are seen here. A certain mass can derive from different structures in other samples, as revealed by LIF MS/MS. The example of *m/z* = 1343 has been discussed above (Figure 5). *m/z* = 1049 in C-5 reminds of the “Raa” glyco-group, but stands for a different structure ^13^. *m/z* = 1269 is a remarkable as in C-46 this glycan was shown to contain fucose ^11^. The new product C-152 exhibited the same MS/MS profile, whereas C-82 differed (Figure S5).

Next, samples were sorted according to their type of Man9 profile (Figure 4). Another sorting criterium was the pattern of high-mass N-glycans, which displayed a startling diversity (Figure 6).

Another notable criterium is the presence of glycans with three or four HexNAc residues, which come with various compositions and apparently do not form a coherent group of structures. Finally, samples may exhibit characteristic peaks of moderate intensity. These are mentioned in a necessarily arbitrary and incomplete manner in Table 1. All spectra are shown in Figure S6.

### 3.8. Towards a single number characterization with PGC-LC-MS

Samples from the same or closely related *Chlorella* strains may - for various reasons - yield MALDI-TOF MS N-glycan patterns with varying peak heights. This “intra-sample” variance reduces the reliability of assignments. LIF-MS/MS spectra, informative as they are, are clumsy and waste space. Here we propose retention-time normalized chromatography on porous graphitic carbon (PGC) as an approach to arrive at numbers that consider “structure” in addition to molecular mass. To make such retentions times useful beyond the narrow scope of this study, they will be expressed as “virtual minutes” (v-min) with the help of isotope-labeled internal standards ^17, 18^. MALDI or ESI based MS/MS spectra can serve to corroborate or falsify an identity insinuated by equal (virtual) retention times. Having served this purpose, they are no longer relevant.

We subjected selected samples containing peaks with *m/z* = 1329 to this PGC-LC-MS regime. One striking result was that the glyco-types Jar, Gov, Asp and even Pit exhibited identical retention times, whereas Sol samples behaved clearly different (Figure 7). Thus, the subtle differences seen in the MALDI-TOF MS spectra, in particular the occurrence of *m/z* = 1460, were based on fundamental structural differences of the complex-type N-glycans of these samples, even though they consist of the same monosaccharide components as recently shown ^11^. The methylated glycans of *m/z* = 1343 eluted after the parent peak, but did not co-elute with that mass from sample C-1 (Figure 7). So, obviously, neither the “Sol” nor the “Jar” and “Gov” glycotypes share the structure found for the C-1 (and C-54, 115, 151 and 108) major glycan.

**Figure 7.**
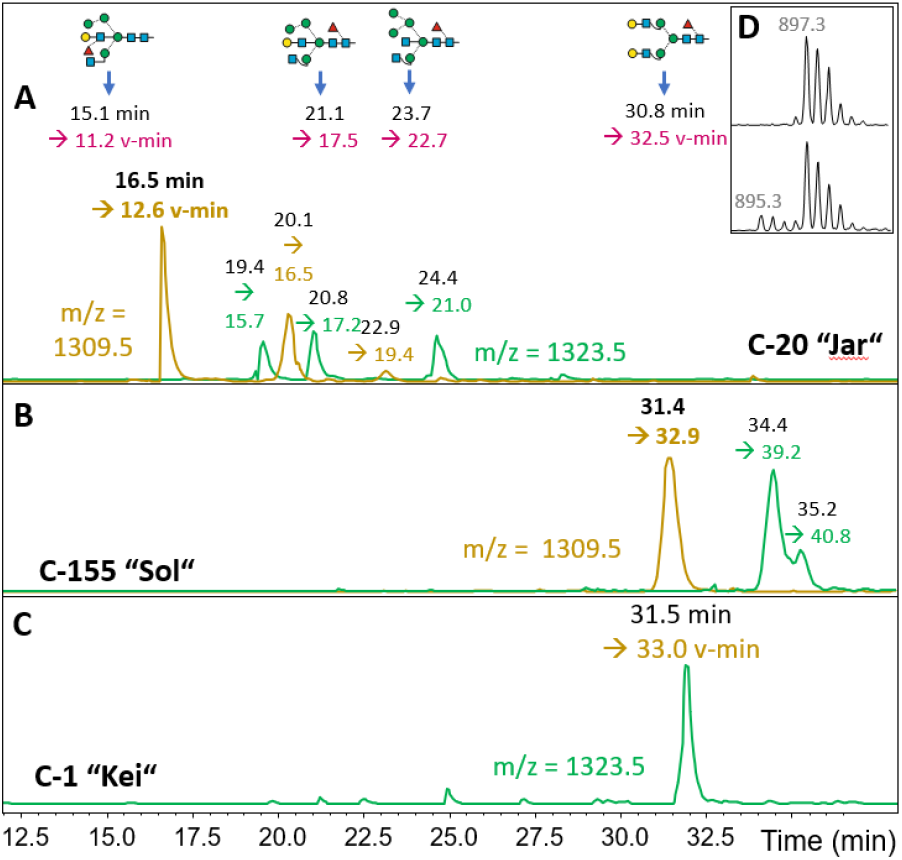
PGC-LC-ESI-MS with conversion to standardized retention times (v-min). Panel **A** shows the extracted ion chromatograms for os3230 and 3231 for a “Jar”-type product (C-20). Panel **B** depicts the elution of os3230 and 3231 in a product with a “Sol”-type MALDI spectrum. These elution characteristics were also found for several other samples (C-15, C25, C-43 for “Jar” and C-21, C-89 and C-125 for “Sol”). Chromatogram **C** shows that the major glycan of a “Kei” sample C-1 does not co-elute with the isobaric glycans of “Jar” and “Sol”. Arrows at the top indicate the elution times of *m/z* = 895.3 brain glycans ^17^ and ^13^C_4_-labeled A^4^A^4^F^6 18^, which were used to calculate v-mins (colored) from the initially measured values (black). Insert **D** shows the spectrum of the isotope-labeled standard A^4^A^4^F^6^ alone or as part of a sample.

Surprising in the contrary sense was the observation that *m/z* = 1329 of C-89, C-125 and C-155 co-eluted with typical Sol samples despite patterns that at first sight lacked an obvious resemblance with the “Sol” glyco-pattern. For consistent results, column temperature needs to be controlled carefully, as the adsorption isotherms of the internal standard and the microalga N-glycan apparently deviated a bit.

As the MALDI-MS/MS patterns of C-5 and the “Raa” products’ *m/z* = 1049 differ only by peak ratios and as the *bona fide* “Raa” products exhibited large differences in the larger glycans (Table 1), examples of each glyco-subtype were subjected to PGC-LC-MS. All *m/z* = 1029 (ESI equivalent to 1049 in MALDI spectra) peaks eluted at 17.3 v-min (Figure S3). Another example for the definition power of PGC-LC-MS was found with sample C-108. The MALDI peak of *m/z* = 1401 in C-108 and in “Hel” samples co-eluted on PGC (Figure S7). It may be noted that this glycan eluted extremely late, *i*.*e*., about 12 min after the glycan A^3^A^3^F^6^, which is the most retained member of the previously used standard series ^17, 19^ indicating identity of this structure. The glycan resembling the major “Kei” structure clearly contained one N-glycan, which co-eluted with the C-1 compound and also gave the same MS/MS spectrum (Figure S7). C-108 contained an additional two peaks with differing spectra (Figure S7), however this observation could be explained.

### 3.8. Contributions and limits of DNA barcoding

The primers used in the recent study fitted to the highly conserved ends of the 18S rRNA and 23S rRNA with optimal homology for *Chlorella*-clade algae ^11^. In eleven cases (C-6, 17, 21, 24, 28, 32, 35, 45, 59 and recently 126) these primers amplified a PCR product and again in many cases this PCR band gave a clear sequence without further subcloning (see “Supplemental sequences in ^11^). Two products (C-36 and C-46) required sub-cloning. In these cases, 15 colonies (out of around 200) were randomly selected for sequencing. For many products, no PCR product was obtained, which might reflect degradation of the DNA or a too low homology of the chosen primers with the sample, which in turn would point at a low kinship of the respective alga with the *Chlorellaceae*.

In case of C-6, C-32, and C-126, where a corresponding live strain was available, the glycan patterns of products and live strains matched almost perfectly. This is a highly important result as it demonstrates a strong link between genome type and N-glycan pattern in products that are derived from one alga strain (scenario A in Figure 6). The other products that gave one barcoding sequence might likewise belong to this group. They may, however, also fall into category C, where two or more algae species constitute the product of which only one is amplified. This concept could explain that many products of the large similarity groups (Raa, Sol, Jar, Gov and Vul) presented with variability in *e*.*g*. the type of methylation of oligomannosidic structures or the occurrence of high-mass glycans.

A brief look at the homologies of the (18S)-ITS1-5.8S-ITS2-(23S) barcodes of strains as well as products supports this notion. The sequence identities between *C. vulgaris* and two different *C. sorokiniana* strains (generating the “Hel” and the “Raa” patterns) are in the range of 75 % (Figure 8). Together with their totally different N-glycan structures (Figure 1), the *C. sorokiniana* strains should probably be considered as two different species. The barcode dissimilarities between *C. vulgaris* and three products likewise indicated different species. Interestingly, there is no higher similarity between *C. vulgaris* and “Gov” on the one hand as insinuated by oligomannose methylation, and between “Jar” and “Gov” on the other hand as suggested by the identical structures of their “pentose-type” glycans (Figure 7). In contrast, “Sol” and “Gov” display a very high sequence identity despite different N-glycan structures (Figure 8).

**Figure 8.**
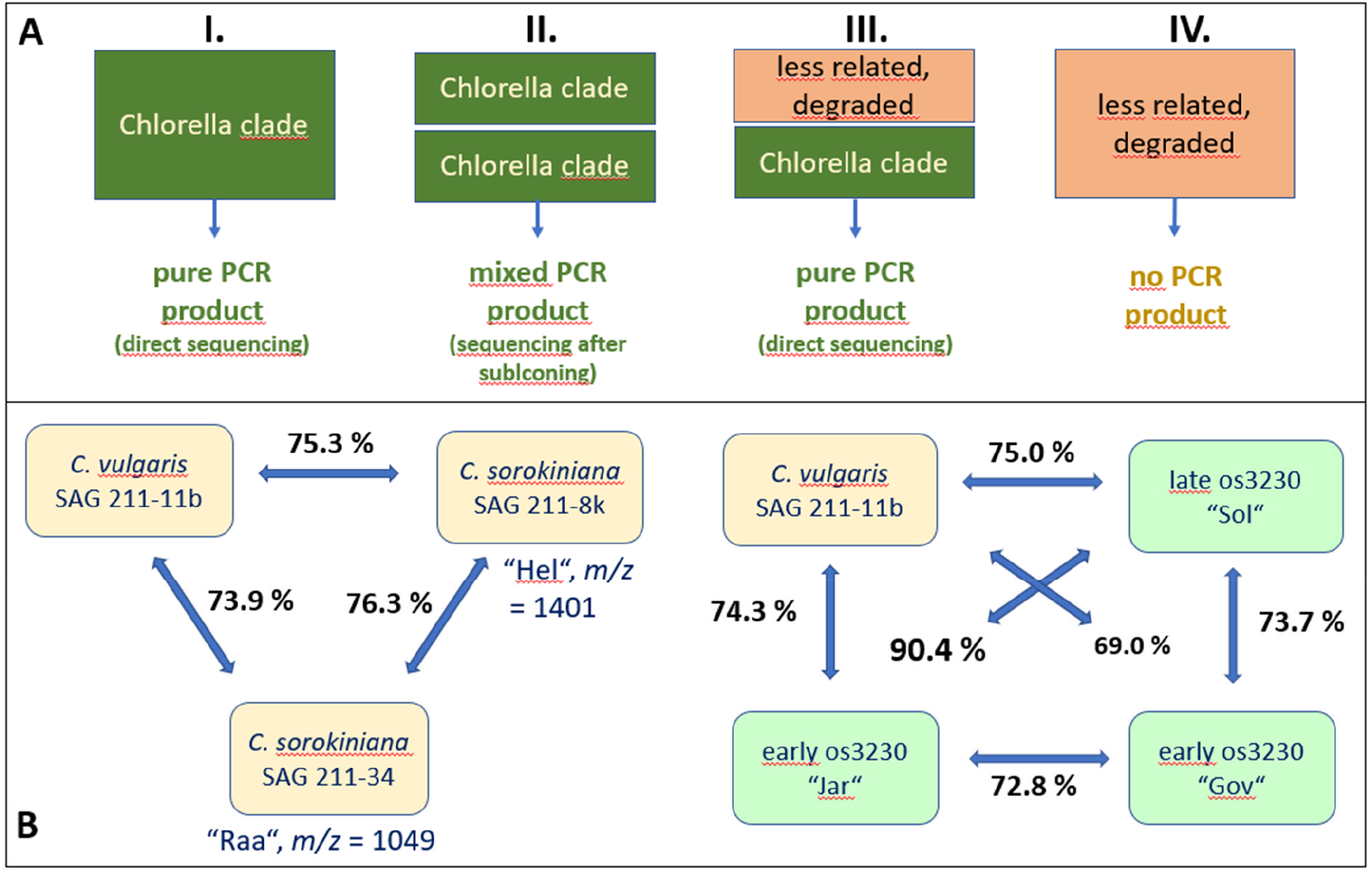
DNA barcoding of Chlorella products. **A**: Scenarios of DNA barcoding with primers designed for *Chlorella*-clade microalgae. A rectangle symbolizes product derived from one microalga strain. **B**: Sequence identities of the ITS1-5.8S-ITS2 region (Supporting Information; Sequences) of selected samples. The left comparison involves strains with known, highly deviating glycan structures. The right block compares strains with partially identical features, *i*.*e*., oligomannose methylation in C. vulgaris and “Gov” on the one and the structure of os3230 (*m/z* = 1329) in “Jar” and “Gov” on the other hand. A complete identity matrix is presented as Figure S7.

With a sequence identity of 96.9 % the products C-1 (“Kei”) and C-23 are close to the point where they could be put into the brackets of one species ^5^. The differing N-glycans yet clearly speak another language (Figure 5). Figure S7 poses more such puzzles that probably await finding of the respective alga strains for explanation.

## DISCUSSION

Analysis of the N-glycans of 172 commercial *Chlorella* samples surprised with a still growing, apparently unlimited variety of N-glycan patterns. At least 72 % of the products fell into a few groups with obvious similarities of their MALDI-TOF MS patterns. MALDI-TOF MS/MS and PGC-LC-MS could in some cases corroborate identities and in other cases demonstrate differences as *e*.*g*., between the “Sol” group and the – at first sight – rather similar supergroup comprising the “Jar”, “Gov” and “Asp” patterns. Some glyco-types may be further splitted (*e*.*g*. the “Raa” group according to the degree of methylation of OM glycans), others may be lumped together, such as the “Jar”, “Gov” and “Asp” groups as they obviously share the same set of medium sized pentose-containing glycans.

A frequent occurrence of a particular glyco-type may arise from a particular strain being cultivated at different sites and times or it may be the result of many vendors obtaining raw material from just a few sources. Retailers are unfortunately hardly ever inclined to reveal the very source of their product. In how far such discretion is appropriate in times of increasing mandate for supply chain transparency could be disputed. This ignorance about the products’ history leaves questions when we look at patterns in which different elements appear to be freely combined, such as just for example the “Raa”-typical *m/z* = 1049 glycan with various types of methylation of oligomannosidic glycans.

Our recent DNA barcoding results revealed that glyco-type based classification by and large matches genetic traits ^11^. The big difference between DNA barcoding and N-glycan profiling lies in the innate ability of glycan profiling to provide a steadfast holistic view of a product’s content of eukaryotic organisms. The presence of a certain glycan structure or glycan pattern is a clear-cut and unambiguous footprint of the production strain(s), whereas sequence differences in a non-functional DNA are a highly nebulous matter. We surmise that base exchanges in a region not subject to selection pressure are a far less solid fundament for classifications than the formation of enzymes of clearly different functionality. Glycan based characterization is independent of the quality of a product’s DNA or of the primers used and will display a more dependable portray of the eukaryotic components of a sample even in case of mixtures. N-glycan patterns or structures are new-comers in the tool-box of microalgae classification. There are already highly sophisticated tools in place for barcode assessments ^4^ and profound considerations on the proper application of DNA barcoding data ^5^. Hesitance towards implementation of yet another aspect is understandable, also in the light of the considerable effort linked to it. Even though, we presume that at least for strains of commercial or intense technical use, stakeholders should not shy away from this effort.

For five glyco-types, the major N-glycans have been structurally elucidated (Figure 1) ^12, 13, 15^. These glycans exhibited a number of features that had not yet been described, neither in plants nor in animal N-glycans. Each of these features requires a specific enzyme for their biosynthesis. Glyco-types differing in several features thus require the emergence of several different glycosyltransferase. The *m/z* = 1401 glycan in “Hel” samples, e.g., requires thee unusual glycosyl- and three specialized methyl-transferases. Even the small *m/z* = 1049 “Raa” glycan could not be formed without three rather individual transferases. The spectra shown in Figures 4, 5 and 6 witness the existence of yet many more unusual transferases. This ingenuity is not restricted to the *Chlorella* clade as demonstrated by a parallel work on *Scenedesmus* N-glycans (R. Mocsai, unpublished data).

Where does this plentitude of glycosyltransferases come from? Eukaryotes and within them green algae are not known to engage mobile genetic elements such as plasmids or, possibly, viruses that might transfer biosynthetic capabilities from one bacterial strain to another. At present, no such mechanism for extra-chromosomal gene transfer in algae would be known. The clear co-clustering of glycosylation traits with rRNA sequences ^11^ rather argues for a chromosomal location of the relevant glycosyltransferases. This, however, would entail that the different glycosyltransferases are the product of evolutionary processes. In appreciation of the fact that land plants were not able to bear a single glyco-enzyme in addition to what mosses already brought in as a founding dowry to the embryophyte taxon, this gain of function leap within what is considered a genus or clade is – cautiously put – remarkable. In other words, we strongly suggest to utilize N-glycan profiles and structures as a relevant element for microalgae classification.

A note of caution is that the herein applied experimental strategy, which involves pepsin digestion at low pH and MALDI-TOF MS of underivatized glycans, is sub-optimal for the detection of sialylated N-glycans. Partial loss of sialic acid and inferior detection of sialylated glycans has been observed with human immunoglobulin and transferrin (data nor shown). However, sialylated species were likewise not detected when preparations were performed at neutral pH and with MALDI-TOF MS analysis of permethylated N-glycans or ESI-MS of glycopeptides did not detect any sialylated species in other microalgae ^20 21 22^.

Back to more practical issues. This work – necessitated by constraints of time, manpower and resources – remains incomplete in many ways. More live strains could be grown and analyzed, more products could be ordered and re-ordered, more LC-MS runs could be performed or more DNA barcoding on more loci could be attempted or alternative genetic analyses could be applied. We think, however, that the value of raising awareness to the just recently discovered possibility of characterizing microalgae strains by their N-glycans rather sooner than later outweighs imperfections that could not be curtailed in a reasonable time span with the resources available. Glycan patterns can serve to identify the origin, identity and purity of microalgal strains and products. While the currently applied procedure requires considerable time and work power, more streamlined processes could be applied to control algae much the same way as *e*.*g*., fish authenticity is analyzed by MALDI-TOF MS ^23^.

It shall be emphasized that by no means do we want to question the presumed health benefits of the products regardless of the particular *Chlorella*-clade strain used. This work shall not at all undermine the reputation of any vendor or producer of algae as all the products can be expected to meet the specifications regarding vitamin and fatty acid content or other quality-determining features. The results presented here should, however, initiate a broad interest in glycan-based strain characterization. We suppose that its application could benefit the food additive market as well as the manifold technical applications of microalgae strains as in both areas the very identity of the used strains seems to be obscured by inappropriate, polysemous species designations.

To conclude, we hope that producers and taxonomists will recognize N-glycans as a valuable if not indispensable element in the toolbox for microalgae classification and identification.

## Abbreviations used

“Hel”, “Raa”, “Sol”, “Jar”, “Gov”, “Asp”, “Vul”, “Kei”: designations of N-glycan patterns;
MALDI-TOF MS: matrix assisted ionization time-of-flight mass spectrometry;
PGC-LC-MS …: liquid chromatography electrospray-ionization mass spectrometry on porous graphitic carbon;
PHexNAc: N-acetylhexosamines;
PNGase A: peptide:N-glycosidase A;
A^4^A^4^F^6^: abbreviation for a biantennary, fucosylated N-glycan according to the proglycan scheme.

## Acknowledgments

This project was supported by the BOKU Core Facility Mass Spectrometry. We are grateful for the excellent technical assistance by Thomas Dalik.

## Supporting Information description

File “Supporting information” containing

Part I: Supporting figures S1 – S7 (except figures S2 and S6)

Part II: Supporting sequences

Part III: Supporting Figure S2 Part IV: Supporting Fig S6

File “Chlorella products Supporting Tables.pdf”

## Footnotes

1 Nominal masses are given for the sake of legibility. Exact masses are found in Figure 3 and Table S4.

