## Supplemental Figures for "The wondrous and worrying diversity of the N‐glycans of *Chlorella* food supplements": 2 Chlorella products Supporting Information.pdf

Department of Chemistry, University of Natural Resources and Life Sciences, Vienna (BOKU)

Part I: Supporting figures (see page 2 ff and the specified Powerpoint files)

Part II: Supporting sequences (see page 7 ff)

Part III: Supporting tables (see Excel file “Chlorella products Supporting Tables“)

### Part I: Supporting figures

**Figure S1** MALDI-TOF MS spectra of live algae strains representing *C. vulgaris* and two different types of *C. sorokiniana.*

***Chlorella vulgaris* strains**

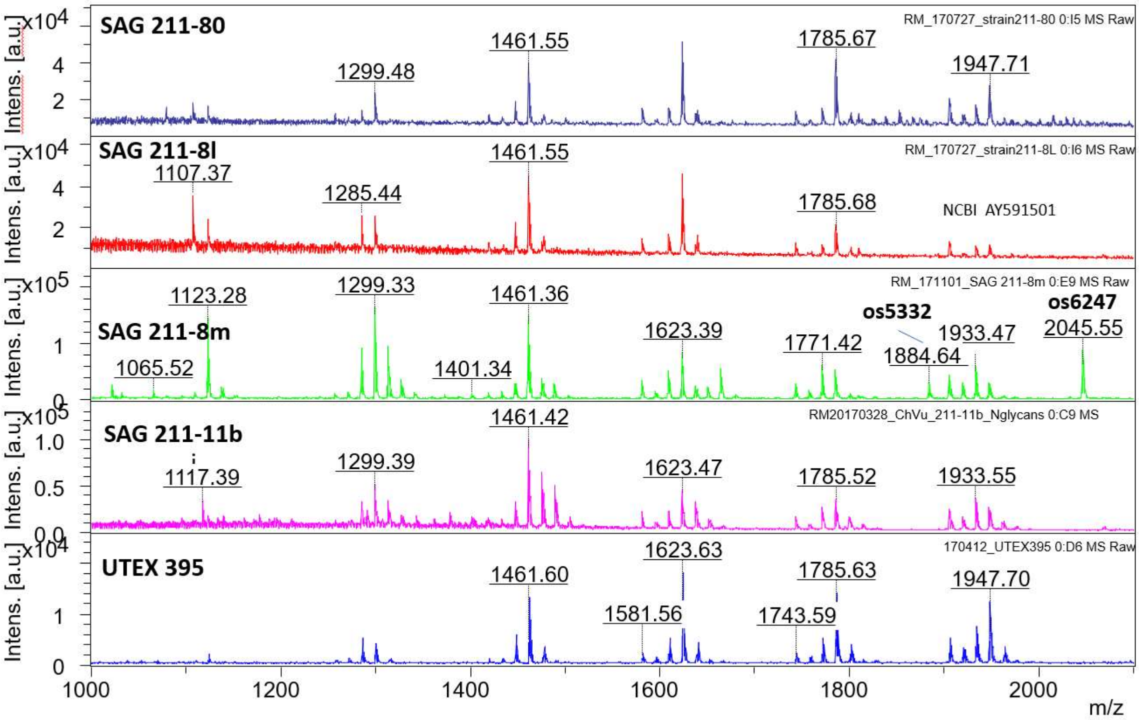


***Chlorella sorokiniana* strains**

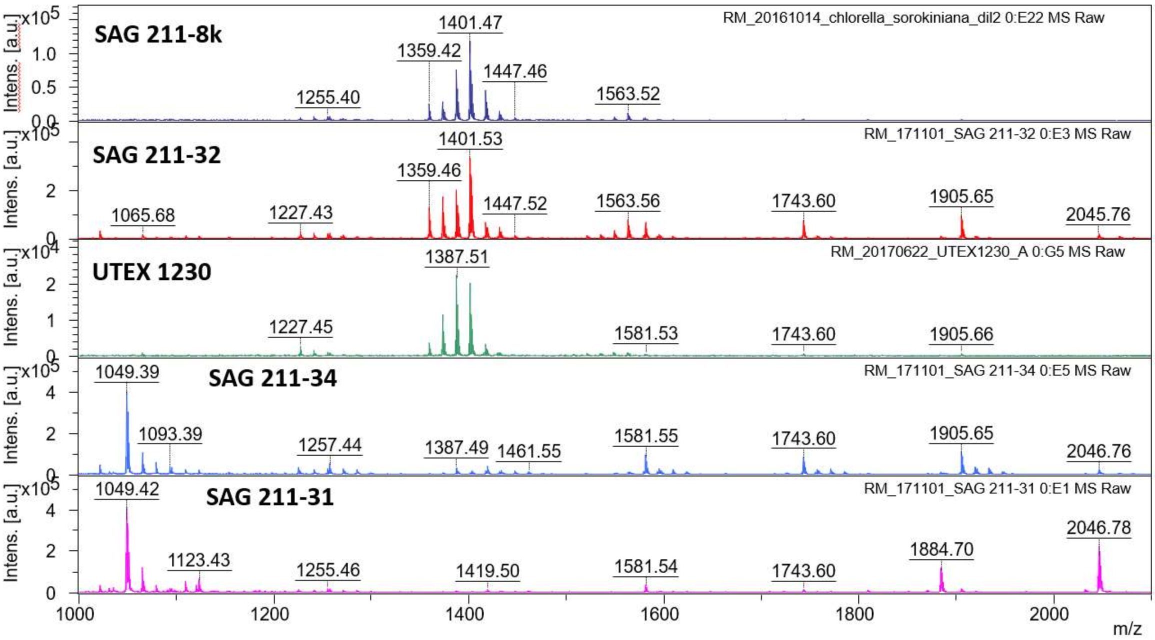

**Figure S2** Exemplary MALDI-TOF MS spectra of samples representing the large N-glycan pattern groups. See extra file “Figure S2 Chlorella products grouped.pdf”

**Figure S3** MALDI-TOF MS/MS spectra of m/z =1049 (os2221) in selected samples. The upper two examples are representative for all samples with a "Raa" glycan pattern irrespective of oligomannose methylation. The bottom spectrum – though similar was derived from a different N-glycan structure {Mocsai, 2021 #450}. Fragment ions are annotated with their numbers of hexose, HexNAc, pentose and methyl constituents. As – in this case – the structures have been determined, fragment cartoons are given here. By the way, this shows that interpretation of fragment ions requires prior knowledge of the structure.

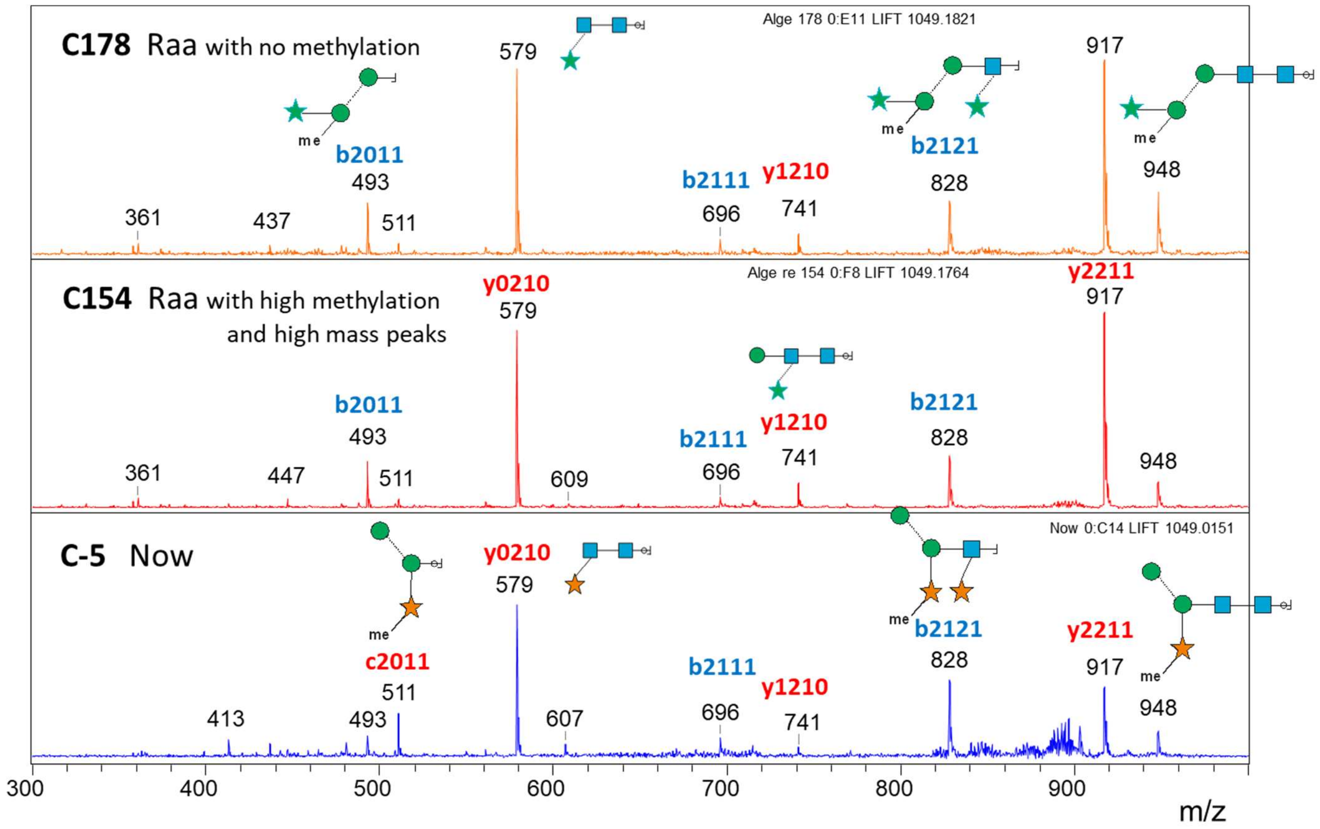

**Figure S4** Scrutiny of os2221 from different "Raa"-type samples by PGC-LC-MS. Panel **A** shows the MALDI-TOF MS spectra of the selected examples. Panel **B** depicts the XICs of the respective $[M+H]^+$ ion together with brain N-glycans that served as internal standard. The peaks with the correct mass are indicated by the blue arrow. All other peaks had differing masses.

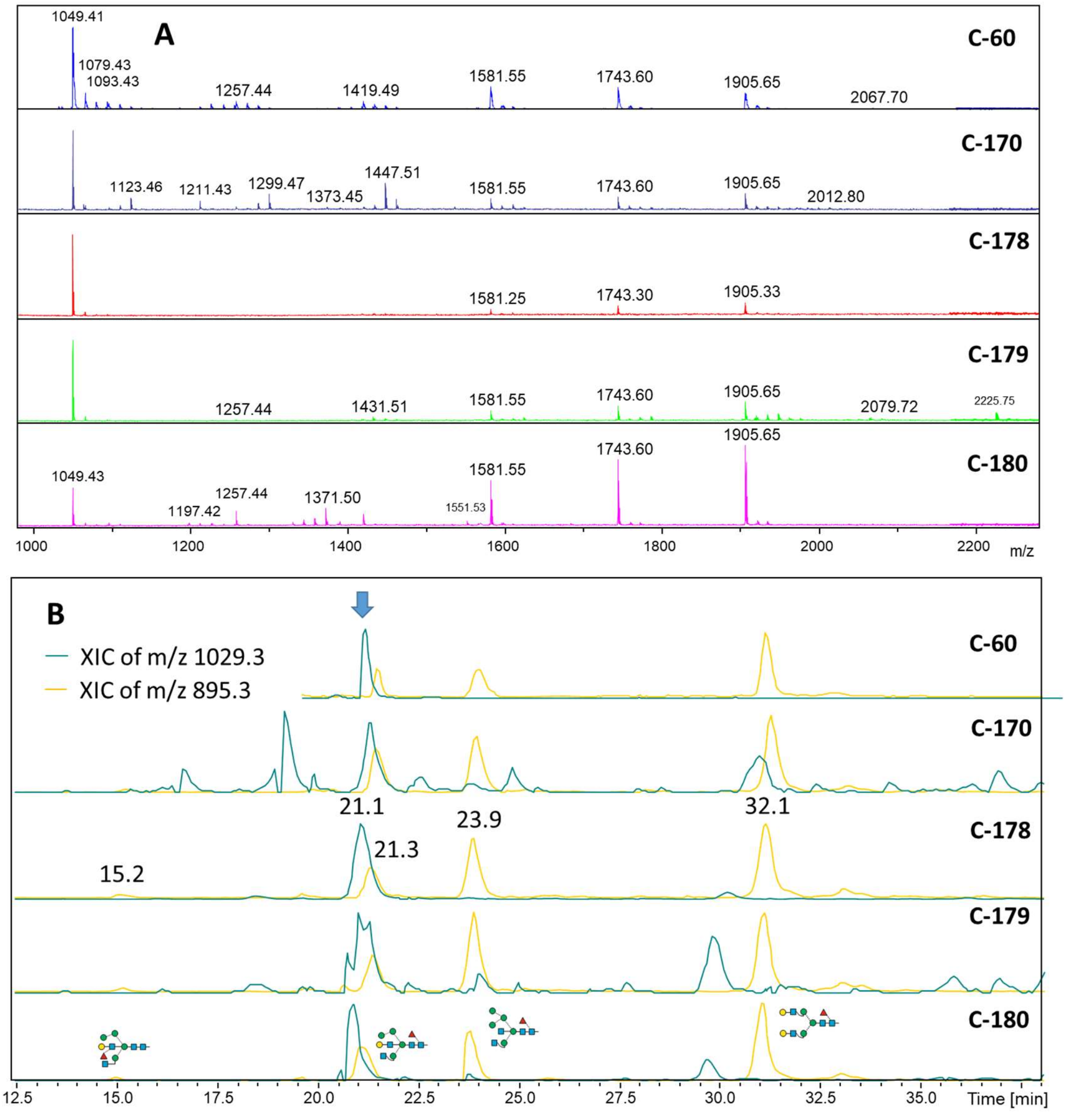

**Figure S5** Comparison of three *m/z* = 1269 peaks by MALDI-TOF MS/MS. Samples C-46 and C-152 apparently contain a substituted (maybe fucose) reducing GlcNAc, while the same peak in C-82 has a clearly different structure.

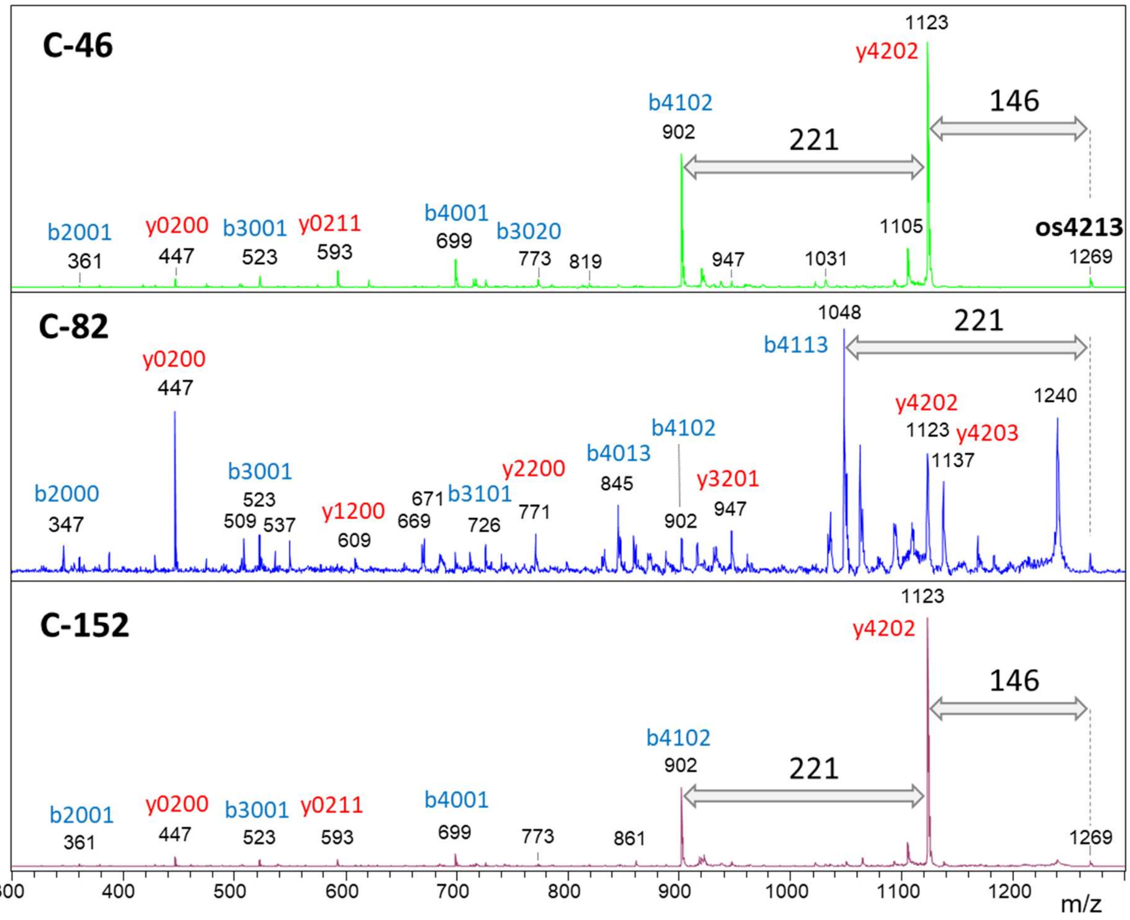

**Figure S6** MALDI-TOF MS spectra of all samples that were not assigned to the large N-glycan pattern groups are shown in the file "Figure S6 Chlorella products spectra unassigned.pdf". The order of these spectra follows that of Table 1.

**Figure S7** Comparison of sample C-108, C-1 and C-32 by MALDI-TOF MS and PGC-LC-MS and MS/MS with emphasis on the MALDI-TOF MS peaks with *m/z* = 1343 (os3231) and 1401 (os4223). Panel **A** gives the MALDI-TOF MS spectra. Panel **B** shows the elution on PGC of os4223 from C-32 and C108 supported by internal standards (brain glycans). Panel **C** shows a similar comparison for os3231 and panel **D** presents the associated MS/MS spectra. Panel **E** demonstrates the difference of spectra for *m/z* = 1323 in sample C-108.

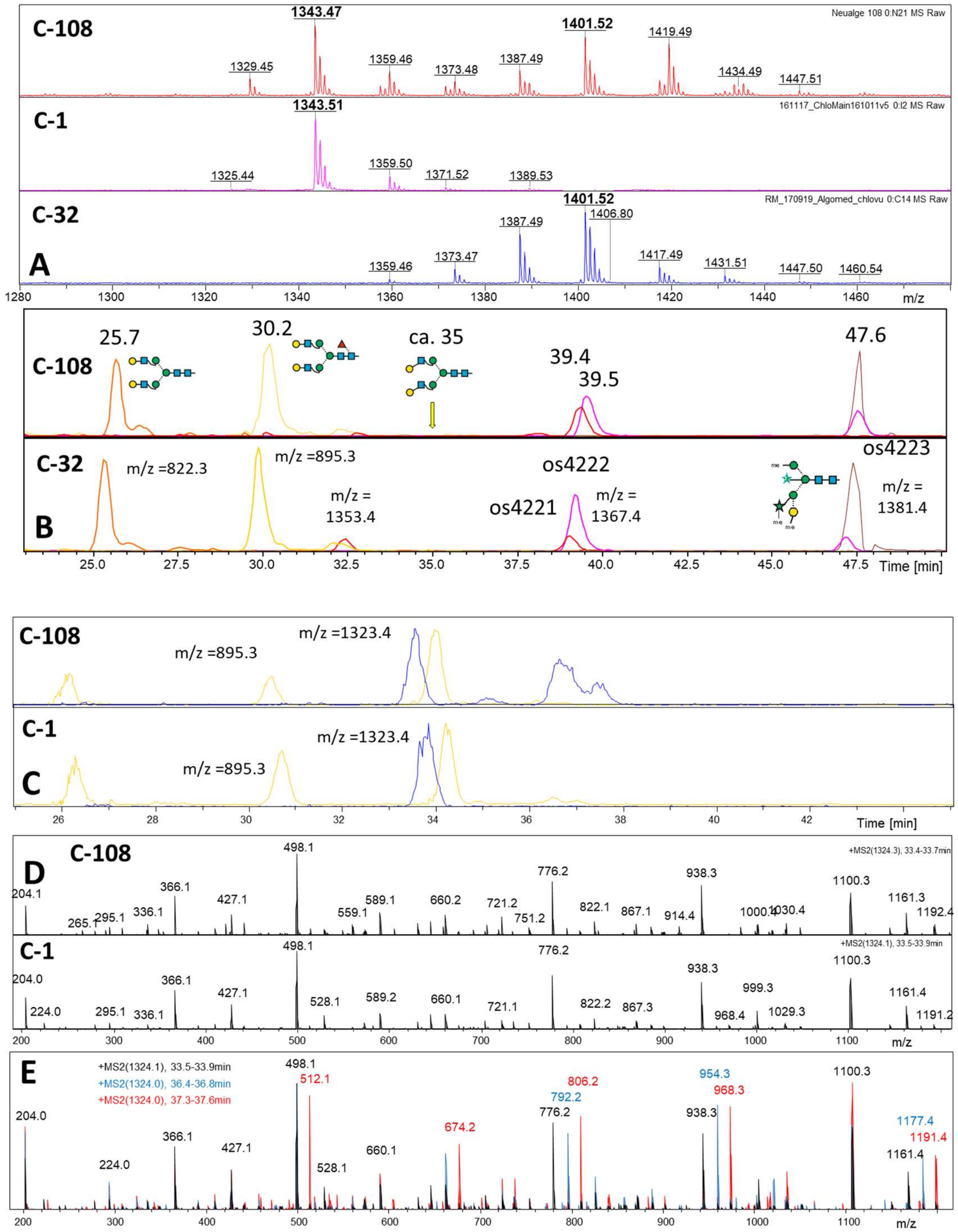

**Figure S8** Identities and deviations of nucleotide sequences of the ITS1-5.8S-ITS2 region as calculated with web.expasy.org/cgi-bin/sim. Considered that a) the more highly conserved 5.8S rDNA accounts of more than a quarter of the sequence and that b) a totally unrelated sequence as *e.g.*, bovine albumin yield “identities” of over 40 %. That means that the yellow and green cells indicate essentially unrelated sequences.

| | **211-11b** | **Hel** | **Raa** | **Sol** | **Jar** | **Gov** | Kei | Jos | Ori28 | Ori46A | Ori46B | Ori46C | SunA | SunB | |
|---|---|---|---|---|---|---|---|---|---|---|---|---|---|---|---|
| | **% identity / Deviation of sequences between TTTCCGTAGGTG... and ...TTCGACCT** | | | | | | | | | | | | | | |
| **211-11b** | 100 | 24.7 | 26.1 | 25 | 25.7 | 31 | 24.6 | 25 | 31.2 | 31.2 | 24.6 | 37.4 | 24.9 | 38.8 | **Deviation** |
| **Hel** | 75.3 | 100 | 23.7 | 9.3 | 9.1 | 28 | 6.1 | 7.3 | 26.8 | 26.2 | 9.8 | 36.4 | 23.7 | 36.8 | |
| **Raa** | 73.9 | 76.3 | 100 | 24.9 | 23.7 | 28.5 | 22.2 | 22 | 27.3 | 26.2 | 23.4 | 36.8 | 16.8 | 37.1 | |
| **Sol** | 75 | 90.7 | 75.1 | 100 | 9.6 | 26.7 | 8.8 | 10 | 26.6 | 26.1 | 8.7 | 36.1 | 24.2 | 36.9 | |
| **Jar** | 74.3 | 90.9 | 76.3 | **90.4** | 100 | 27.2 | 6.1 | 5.7 | 25.9 | 25.7 | 10.9 | 36.4 | 23 | 36.7 | |
| **Gov** | 69 | 72 | 71.5 | **73.3** | **72.8** | 100 | 27.7 | 28.8 | 15.8 | 14.8 | 26.6 | 34.1 | 26.8 | 35.3 | |
| Kei | 75.4 | 93.9 | 77.8 | 91.2 | 93.9 | 72.3 | 100 | 3.1 | 26.1 | 26.2 | 9.7 | 35.1 | 22.5 | 36.2 | |
| Jos | 75 | 92.7 | 78 | 90 | 94.3 | 71.2 | 96.9 | 100 | 26.2 | 25.8 | 10.7 | 35.4 | 22 | 36.9 | |
| Ori28 | 68.8 | 73.2 | 72.7 | 73.4 | 74.1 | 84.2 | 73.9 | 73.8 | 100 | 16 | 25.3 | 35.8 | 25.4 | 37.5 | |
| Ori46A | 68.8 | 73.8 | 73.8 | 73.9 | 74.3 | 85.2 | 73.8 | 74.2 | 84 | 100 | 25.6 | 35.5 | 24.6 | 35.9 | |
| Ori46B | 75.4 | 90.2 | 76.6 | 91.3 | 89.1 | 73.4 | 90.3 | 89.3 | 74.7 | 74.4 | 100 | 35.8 | 25.4 | 36.5 | |
| Ori46C | 62.6 | 63.6 | 63.2 | 63.9 | 63.6 | 65.9 | 64.9 | 64.6 | 64.2 | 64.5 | 64.2 | 100 | 35 | 3.9 | |
| SunA | 75.1 | 76.3 | 83.2 | 75.8 | 77 | 73.2 | 77.5 | 78 | 74.6 | 75.4 | 74.6 | 65 | 100 | 35.3 | |
| SunB | 61.2 | 63.2 | 62.9 | 63.1 | 63.3 | 64.7 | 63.8 | 63.1 | 62.5 | 64.1 | 63.5 | 96.1 | 64.7 | 100 | |
| | **Identity** | | | | | | | | | | | | | | |

#### Part II: Supporting sequences

**Part IIa**: Sequences from Mocsai *et al.* 2019 used for identity assessment.

Nucleotide sequences of ITS1-5.8S-ITS2 rRNA gene with flanking regions of 18S and 26S rDNA. Identity assessment using web.expasy.org/sim/ was performed without the highly homologous regions at the 5' and 3' end (in brackets). A highly conserved region comprising 23 % of the compared sequence – roughly representing the 5.8S rRNA - is underlined. Most of the observed divergences reside in the regions before (ITS1) and after (ITG2) this 5.8S sequence.

>Kei_C1

(ACACACCGCCCGTCGCTCCTACCGATTGGGTGTGCTGGTGAAGTGTTCGGATTGGCGACCGGGGGCGGTCTCCGCTCTCGGCCGCCGAGAAGTTCATTAAACC
CTCCCACCTAGAGGAAGGAGAAGTCGTAACAAGG)
TTTCCGTAGGTGAACCTGCGGAAGGATCATTGAATCGATCGAATCCACACCGGTAACCACACGTCGCCCCTGTGGTGCATTCGCCGACCTCCGGCGTTTCACCCT
GGCGTCGGCCCCTGGGCTGGGGCTCTCACGAGCCGCTTTCCAGGTCCGACGGGCGCCTCCCTTGGGCTCACCCCCTGGGGCTGGCGTCGGCCAAAACCCCTGTA
TCCAACCCTTTTTTTAACACACCCCAAACCACAACCAACTCTGAAGCATCTTTGGTGGCCCGGCCCCGTGCCGTCCACTCCAAACCAAAGACAACTCTCAACAACG
GATATCTTGGCTCCCGTATCGATGAAGAACGCAGCGAAATGCGATACGTAGTGTGAATTGCAGAATTCCGTGAACCATCGAATCTTTGAACGCAAATTGCGCCC
GAGGCTTCGGCCGAGGGCATGTCTGCCTCAGCGTCGGTTTACACCCTCGCCCTCCCCCACCCTGTGTGGTGTGGTTGGTGCGGATCTGGCCCTCCCGGCTCCGCT
CTCCTTGAGCGTCCGGGTTGGCTGAAGTGGAGAGGCTTGAGCATGGACCCCGTTTGTAGGGCAATGGCTTGGTAGGTAGGCACCCCCTACGCAGCCTGCCGTT
GCCCGAGGGGACTTTGCTGGAGGCCCAGCAGGAATCCGGCTGTCTTTGGCAGCCGGACTACTCACTCATTCGACCT
(GAGCTCAGGCAAGA)

>Hel_C32

(ACACACCGCCCGTCGCTCCTACCGATTGGGTGTGCTGGTGAAGTGTTCGGATTGGCGACCGGGGGCGGTCTCCGCTCTCGGCCGCCGAGAAGTTCATTAAACC
CTCCCACCTAGAGGAAGGAGAAGTCGTAACAAGG)
TTTCCGTAGGTGAACCTGCGGAAGGATCATTGAATCGATCGAATCCACACCGGTAACCACACTGTCGCCCTCGGCGGTGCATTCTCTGGCTTCGGCTGGGTTTCA
CCCCGAGCGTCGGCCCCTGGGTTGGGGTTCTCACGAGCCGCTCTCCAGGTCCGGCGGGCGCCTCCCTTGGGCTCACCCCCTGGGGCTGTCGTCGGCCAAAACCC
CTGTATCCAACCTTTTTTTTAACACACCCCAAACCACAACCAACTCTGAAGCATCTTTGGTGGCCCGGCCCCGTGCCGTCCACTCCAAACCAAAGACAACTCTCAA
CAACGGATATCTTGGCTCCCGTATCGATGAAGAACGCAGCGAAATGCGATACGTAGTGTGAATTGCAGAATTCCGTGAACCATCGAATCTTTGAACGCAAATTG
CGCCCGAGGCTTCGGCCGAGGGCATGTCTGCCTCAGCGTCGGTTTACACCCTCGCCCTCCCCCCCTGTGGGGGGCGGTGCGGACCTGGCCCTCCCGGCTCCGCT
CTCTCCCGAGCGTCCGGGTTGGCTGAAGCACAGAGGCTTGAGCATGGACCCCGTTTGTAGGGCAATGGCTTGGTAGGTAGGCACCCCCTACGCAGCCTGCCGT
TGCCCGAGGGGACTTTGCTGGAGGCCCAGCAGGAATCCGGCCCTTCCCGGCCGGACTACTCACTCATTCGACCT
(GAGCTCAGGCAAGA)

>Raa_C6

(ACACACCGCCCGTCGCTCCTACCGATTGGGTGTGCTGGTGAAGTGTTCGGATTGGCGACCGGGTGCGGTCTCCGCTCTCGGCCGCCGAGAAGTTCATTAAACC
CTCCCACCTAGAGGAAGGAGAAGTCGTAACAAGG)
TTTCCGTAGGTGAACCTGCGGAAGGATCATTGAATCGATCGAATCCACTCTGTGAACCAAACGTCCCCCCTTGGGTGCGGGCTTCGGTCTGCCCCAAGGCGTCG
GTTCCCTGGCTGGGGTCTTCGGACCGCAGTTAGGTCCGGCGGGCGCGCCCTCTGGCGTGTCGGCCCTCGTGGCTGCCGCCAGTTGGGTTCGCTGGAAATTGTAT
CCAACTCAACCCACCCCAAACCACAACTTATACTGAAGCAATCGGTGAGTGCACTCTGGTGCCTCGCTCTAACCAAAGACAACTCTCAACAACGGATATCTTGGC
TCCCGTATCGATGAAGAACGCAGCGAAATGCGATACGTAGTGTGAATTGCAGAATTCCGTGAACCATCGAATCTTTGAACGCAAATTGCGCCCAAGGCTTCGGC
CGAGGGCATGTCTGCCTCAGCGTCGGCTTACCCCCTCGCTCCCCCTCTCCTTTGGAGTGGGTGAACGGATCTGGTTTTCCCGGCTACGTGCTTCTGCACGCCCGG
GTTGACTGAAGTGTAGAGGCTTGAGCATGGACCCCGTTTGTAGGGCAATGGCTTGGTAGGTAGCTTAGCTACACCGCCTGCCGTGGTCCGAGGGGACTTTGCT
GGCGGCCCAGCAGGAATTCGGGTGTTGGGTTTCCCACCCCGAAAGCTTCAAACCTTCGACCT
(GAGCTCAGGCAAGA)

>Sol_C21

(ACACACCGCCCGTCGCTCCTACCGATTGGGTGTGCTGGTGAAGTGTTCGGATTGGCGACCGGGGGCGGTCTCCGCTCTCGGCCGCCGAGAAGTTCATTAAACC
CTCCCACCTAGAGGAAGGAGAAGTCGTAACAAGG)
TTTCCGTAGGTGAACCTGCGGAAGGATCATTGAATCGATCGAATCCACACCGGTAACCACACGTCGCCCTGTGCGGTGCTGCACTCAGCCGAGTGCACTCTGCG
CAGCGTCGGCCCCTGGGCTGGGGCTCTCACGAGCCGCTTTCCAGGTCCGACGGGCGCCTCCCTTGGGCTCACCCCCCGGGGCTGTCGTCGGCCAAAACCCCTGT

ATCCAACCCCTTTTTTTAACACACCCCAAACCACAACCAACTCTGAAGCATCTTTGGTGGCCCGGCCCCGTGCCGTCCACTCCAAACCAAAGACAACTCTCAACAA
CGGATATCTTGGCTCCCGTATCGATGAAGAACGCAGCGAAATGCGATACGTAGTGTGAATTGCAGAATTCCGTGAACCATCGAATCTTTGAACGCAAATTGCGC
CCGAGGCTTCGGCCGAGGGCATGTCTGCCTCAGCGTCGGTTTACACCCTCGCCCTCCCCCACCCTGTGCGGTGGGGTGCAGGTGCGGACCTGGCCCTCCCGGCT
CCGCCCCGCTGTTCTTCGAGCAGCGGTGGCGCCCGGGTTGGCTGAAGCACAGAGGCTTGAGCATGGACCCCGTTTGTAGGGCAATGGCTTGGTAGGTAGGCAC
CCCCTACGCAGCCTGCCGTTGCCCGAGGGGACTTTGCTGGAGGCCCCGCAGGAATCCGGCCCGCCTTTGCGGGCGGCCGGAGCACTCACTCATTCGACCT
(GAGCTCAGGCAAGA)

#### >Jar_C45

(ACACACCGCCCGTCGCTCCTACCGATTGGGTGTGCTGGTGAAGTGTTCGGATTGGCGACCGGGGGCGGTCTCCGCTCTCGGCCGCCGAGAAGTTCATTAAACC
CTCCCACCTAGAGGAAGGAGAAGTCGTAACAAGG)
TTTCCGTAGGTGAACCTGCGGAAGGATCATTGAATCGATCGAATCCACACCGGTAACCAAACGTCGCCCCCCTGTGGTGCATTCGCCGGACCCCCGGCGTTTCA
CCCTGGGCGTCGGCCCCTGGGCTGGGGCTCTCACGAGCCGCTTCTCAGGTCCGACGGGCGCCTCCCTTGGGCTCACCCCCCGGGGCTGGCGTCGGCCAAAACC
CCTGTATCCAACCCTTTTTTAACACACCCCAAACCACAACTCACTCTGAAGCATCTTTGGTGGTCGTGCCTCGTGCCCTTCCACTCCAAACCAAAGACAACTCTCAA
CAACGGATATCTTGGCTCCCGTATCGATGAAGAACGCAGCGAAATGCGATACGTAGTGTGAATTGCAGAATTCCGTGAACCATCGAATCTTTGAACGCAAATTG
CGCCCGAGGCTTCGGCCGAGGGCATGTCTGCCTCAGCGTCGGTTTACACCCTCGCCCTCCCCCACCCTGCGTGGTGGGGTGCTGGTGCGGATCTGGCCCTCCCG
GCTCCCCTCTCCCTCCCAGGCGAGGCTCCGGGTTGGCTGAAGCACAGAGGCTTGAGCATGGACCCCGTTTGTAGGGCAATGGCTTGGTAGGTAGGCACCCCCTA
CGCAGCCTGCCGTTGCCCGAGGGGACTTTGCTGGAGGCCCAGCAGGAATCCGGTCGGTCCCTGTGGCCGCCGGACCACTCACTCATCGACCT

#### >Gov_C35

(ACACACCGCCCGTCGCTCCTACCGATTGGGTGTGCTGGTGAAGTGTTCGGATTGGCGACCCGGGGCGGTTTCCGCCCTGGGCTGCCGAGAAGTTCATTAAACC
CTCCCACCTAGAGGAAGGAGAAGTCGTAACAAGG)
TTTCCGTAGGTGAACCTGCGGAAGGATCATTGAATCGATCGAATCCACACCGGTAACCATCCTACCCCCCCTGGCCTAACACCCCAGGGGCGCCAGTCCCCTGG
CCCGGGCCCGACCCGGTGCCCAGGTCTGGCGGGGTGGGGGCCCGTCCCCCGCCTGGTAATTTGTCCAACCTTAACACACCCCAAACGTCAAAACCAAACTGAAG
CAACTGGACTGGGCGGCCCCAGCGCCCCCCATCCGCCAAACCAAAGACAACTCTCAACAACGGATATCTTGGCTCCCGTATCGATGAAGAACGCAGCGAAATGC
GATACGTAGTGTGAATTGCAGAATTCCGTGAACCATCGAATCTTTGAACGCAAATTGCGCCCGCGGCTCCGGCCAAGGGCATGCCTGCCTCAGCGTCGGCTTTC
ACCCCCTCGCCCCCAATACATTTGGGAGCGGACCTGGCACCCTCGGGGGCCCGGCCTTTTCCAAAGGCCGCCGCCCCCGGGCCTGCTGAAGTGCAGTGGCTTGA
GCATGGACCCCGTTTGCAGGGCAATGGCTTGGTAGGTAGGCCCCGGCCTGCACCCCGCCTGCCGTTGCCTGAGGGGACTTTGCTGGGAGCCTAGCAGGAATTG
GGAGCCCCAGCCCTGGCGCTGGGCCCCAACCCCCCCCATATTTCGACCT (GAGCTCAGGCAAGG)

#### >Asp_C59

(ACACACCGCCCGTCGCTCCTACCGATTGGGTGTGCTGGTGAAGTGTTCGGATTGGCGACCCGGGGCGGTTTCCGCCCTGGGCTGCCGAGAAGTTCATTAAACC
CTCCCACCTAGAGGAAGGAGAAGTCGTAACAAGG)
TTTCCGTAGGTGAACCTGCGGAAGGATCATTGAATCGATCGAATCCACACCGGTAACCATCCTACCCCCCCTGGCCTAACACCCCAGGGGCGCCAGTCCCCTGG
CCCGGGCCCGACCCGGTGCCCAGGTCTGGCGGGGTGGGGGCCCGTCCCCCGCCTGGTAATTTGTCCAAGCTTAACACACCCCAAACGTCAAAACCAAACTGAA
GCAACTGGACTGGGCGGCCCCAGCGCCCCCCATCCGCCAAACCAAAGACAACTCTCAACAACGGATATCTTGGCTCCCGTATCGATGAAGAACGCAGCGAAAT
GCGATACGTAGTGTGAATTGCAGAATTCCGTGAACCATCGAATCTTTGAACGCAAATTGCGCCCGCGGCTCCGGCCAAGGGCATGCCTGCCTCAGCGTCGGCTT
TCACCCCCTCGCCCCCAATACATTTGGGAGCGGACCTGGCACCCTCGGGGGCCCGGCCTTTTCCAAAGGCCGCCGCCCCCGGGCCTGCTGAAGTGCAGTGGCTT
GAGCATGGACCCCGTTTGCAGGGCAATGGCTTGCTAGGTAGGCCCCGGCCTGCACCCCGCCTGCCGTTGCCTGAGGGGACTTTGCTGGGAGCCTAGCAGGAAT
TGGGAGCCCCAGCCCTGGCGCTGGGCCCCAACCCCCCCCATATTTCGACCT (GAGCTCAGGCAAGG)

#### >Jos_C24

(ACACACCGCCCGTCGCTCCTACCGATTGGGTGTGCTGGTGAAGTGTTCGGATTGGCGACCGGGGGCGGTCTCCGCTCTCGGCCGCCGAGAAGTTCATTAAACC
CTCCCACCTAGAGGAAGGAGAAGTCGTAACAAGG)
TTTCCGTAGGTGAACCTGCGGAAGGATCATTGAATCGATCGAATCCACACCGGTAACCAAACGTCGCCCCCCTGTGGTGCATTCTCCGGACATCCGGCGTTTCAC
CCTGGGCGTCGGCCCCTGGGCTGGGGCTCTCACGAGCCGCTTTCTAGGTCCGACGGGCGCCTCCCTTGGGCTCACCCCCCGGGGCTGGCGTCGGCCAAAACCCC
TGTATCCAACCCTTTTTTAACACACCCCAAACCACAACCAACTCTGAAGCATCTTTGGTGGTCCGGCCTCGTGCCGTCCACTTCAAACCAAAGACAACTCTCAACA
ACGGATATCTTGGCTCCCGTATCGATGAAGAACGCAGCGAAATGCGATACGTAGTGTGAATTGCAGAATTCCGTGAACCATCGAATCTTTGAACGCAAATTGCG
CCCGAGGCTTCGGCCGAGGGCATGTCTGCCTCAGCGTCGGTTTACACCCTCGCCCTCCCCCACCCTGTGTGGTGGGGTGTTGGTGCGGATCTGGCCCTCCCGGC
TCCGCTCTGATGAGCGTCCGGGTTGGCTGAAGTGCAGAGGCTTGAGCATGGACCCCGTTTGTAGGGCAATGGCTTGGTAGGTAGGCACCCCCTACGCAGCCTG
CCGTTGCCCGAGGGGACTTTGCTGGAGGCCCAGCAGGAATCCGGCTGTTTCGGCAGCCGGACTACTCACTCATTCGACCT (GAGCTCAGGCAAGA)

>Sun_C36_A_abundant_clone

(ACACACCGCCCGTCGCTCCTACCGATTGGGTGTGCTGGTGAAGTGTTCGGATTGGCATCTGGGGGCGGTCTCCGCTTCCTGACGCCGAGAAGTTCATTAAACCC
TCCCACCTAGAGGAAGGAGAAGTCGTAACAAGG)
TTTCCGTAGGTGAACCTGCGGAAGGATCATTGAATCGATCGAATCCACTCTGGTAACCAAACGTCCCCCCCTTGGTGGCAGGGCTTGCCTTGTCCCATGGGCGCC
GGTCCCCTGGCTGGGGCCTTCGGGCCGCAGTTAGGTCCGGCGGGTGTCCCTCCGATGCTGGGGCTTTTGCCCCTCTTCGGTTGGTGATGCTGGAAATTTATATTC
AACTCAACCCACCCCAAACCTCGAATTAATCTGAAGCTGTCTTGTGTCACGCCTCGGCGTAGCACTCTAACCAAAGACAACTCTCAACAACGGATATCTTGGCTCC
CGTATCGATGAAGAACGCAGCGAAATGCGATACGTAGTGTGAATTGCAGAATTCCGTGAACCATCGAATCTTTGAACGCAAATTGCGCCCAAGGCTTCGGCCAA
GGGCATGTCTGCCTCAGCGTCGGCTTACCCCCTCACCCTCCCAATCCCTGTGATTGGGCAGAGTGGATCTGGCCCTCCCGGCTCCGTTCCAACTTGTTGGCACGC
CCGGGTCGGCTGAAGTGTAGAGGCTTGAGCATGGACCCCGTTTGTAGGGCAATGGCTTGGTAGGTAGCCTCTGGTTACATCGCCTGCCGTTGTCCGAGGGGAC
TTTGCTGGCGGCCCAGCAGGAATTTGGTGCGTGCGGTTCTCCGTCGCCCAAATGCTTCACACCTTCGACCT (GAGCTCAGGCAAGA)

> Sun_C36_B_rare_clone

(ACACACCGCCCGTCGCTCCTACCGATTGGGTGTGCTGGTGAAGTGTTCGGATTGGCAGCTTAGGGTGGCAACACCTCAGGTCTGCCGAGAAGTTCATTAAACC
CTCCCACCTAGAGGAAGGAGAAGTCGTAACAAGG)
TCTCCGTAGGTGAACCTGCGGAGGGATCATTGAATTATTAAAACCACAATGTGAACCTCAACGTTCCGTGCCCTGGCTTGCCAGTGGGGCGACATGGTCAACAC
CAGGTCGTACTCACAGCTGGGTGGGCATTGTTGCCTACTCAGTGGCGCCTTGGCATGATCATACACCAGTGCTAACCACTGATAAAACCAAACTCTGAAGTTTGA
TTGCTATTCATTGGCAATCTTAACCAAAGACAACTCTCAACAACGGATATCTTGGCTCTCGCAACGATGAAGAACGCAGCGAAATGCGATACGTAGTGTGAATTG
CAGAATTCCGTGAACCATCGAATCTTTGAACGCATATTGCGCTCGAGCCTTCGGGCAAGAGCATGTCTGCCTCAGCGTCGGTTTAATCCCTCACCCCTCCCTATTA
TGGGTGCGTTGATCATGTGATCAGCCATTGGGGTGGATCTGGCTTCCCCAATCTCACTTGTTGCGATTGGGTTGGCTGAAGCACAGAGGCTTAAGCAAGGACCC
GATATGGGCTTCAACTGGATAGGTAGCAACGGCGTATGCCGACTACACGAAGTTGTTGCTTGTGGACTTTGTTAGGAGCCGAGCAGGAACATGCCTTGTGCAT
GCCTAAACTTTCGACCT (GAGCTCAGGCAAGG)

>Ori_C28

(ACACACCGCCCGTCGCTCCTACCGATTGGGTGTGCTGGTGAAGTGTTCGGATTGGCAGCCCGGGGCGGTTCCCGCTCTGGTTTGCCGAGAAGTTCATTAAACCC
TCCCACCTAGAGGAAGGAGAAGTCGTAACAAGG)

TTTCCGTAGGTGAACCTGCGGAAGGATCATTGAATCGATCGAACCCACACCGGTAACCACACAACCCCCCCTGGCGGCACGCCCCAGGGGCGCCAGTCCCCTGG
CCCGGGCCACAACCCCGGTGCCCAGGTCTGGCGGGGTGTGCCCAGCCCCCGGGCTGGGCACGCCTGGTAATTCTGTCCAACCTCAACCCATCCCAAACCCCAAA
CCAAACTGAAGCTCGACTGGAAGGGCGGCTCTCAGCAGCCCCCGACCACAAACCAAAGACAACTCTCAACAACGGATATCTTGGCTCCCGTATCGATGAAGAA
CGCAGCGAAATGCGATACGTAGTGTGAATTGCAGAATTCCGTGAACCATCGAATCTTTGAACGCAAATTGCGCCCGAGGCTCCGGCCAAGGGCATGCCTGCCTC
AGCGTCGGCTCACACCCCTTGCCCCCCCCACCTGCTGGGGGGAGCAGACCTGGCACCCTCGGGCCAGCCTGGATTGGCTCTCCAGTCCAGCTGTGCCCGGGCCT
GCTGAAGTGCAGAGGCTTGAGCATGGACCCCGTTTGCAGGGCAATGGCTTGGTAGGCTGGCCTTCACGGCTGAGCACCGCCTGCCGTTGCCTGAGGGGACTTT
GCTGGGAGCCCAGCAGGAATTGGGGGCAGCCCTCACCGGCCCCCCAACCCTCTCACTTCGACCT (GAGCTCAGGCAAGA)

>Ori_C46_A

(ACACACCGCCCGTCGCTCCTACCGATTGGGTGTGCTGGTGAAGTGTTCGGATTGGCAACCGGGGGCGGTCTCCGCTCCGGGTTGCTGAGAAGTTCATTAAACC
CTCCCACCTAGAGGAAGGAGAAGTCGTAACAAGG)
TTTCCGTAGGTGAACCTGCGGAAGGATCATTGAATCGATCGAATCCACACCGGTAACCAACCTACCCCCCCTGGCCTCAACCCCCAGGGGCGCCAGTCCCCTGG
CCCCGGCCCCCTGCCCCGTGCAGGGCCCGGGTGCCCAGGTCTGGCGGGGTGGCCCTCGGGCTGCCTGGTAATTGTCCAACCTCAACACACCCCAAACACCTAAC
CACACTGAAGCAATCGGAGCGGCGGCCTCGGCCCCCAATCCACAAACCAAAGACAACTCTCAACAACGGATATCTTGGCTCCCGTATCGATGAAGAACGCAGC
GAAATGCGATACGTAGTGTGAATTGCAGAATTCCGTGAACCATCGAATCTTTGAACGCAACTTGCGCCCGCGGCTCCGGCCAAGGGCATGTCTGCCTCAGCGTC
GGCACACACCCCTCGCCCCCCCCACCGGGTGGGGAGTGGACCTGGCACCCCCAGGCCTCGGCCAGCCCTCACCGGCTGTCGCTGGCCTGGGTCTGCTGAAGTG
CAGAGGCTTGAGCATGGACCCCGTTTGCAGGGCAATGGCTTGGTAGGTAGGCGCCAGCCTGCACCCCGCCTGCCGTTGCCTGAGGGGACTTTGCTGGGAGCCC
AGCAGGAATTGGGGCCCGCCCCGGCGGCCCCAACCCCTTCTCACTTCGACCT (GAGCTCAGGCAAGA)

>Ori_C46_B

(ACACACCGCCCGTCGCTCCTACCGATTGGGTGTGCTGGTGAAGTGTTCGGATTGGCGACCGGGGGCGGTCTCCGCTCTCGGCCGCCGAGAAGTTCATTAAACC
CTCCCACCTAGAGGAAGGAGAAGTCGTAACAAGG)
TTTCCGTAGGTGAACCTGCGGAAGGATCATTGAATCGATCGAATCCACACCGGTAACCACACTGTCGCCCTGGGTCGGGTGCGCACCTCTGCGTGCTGCCCGGC
CCAGCGCCGGCCCCTGGGCTGGGGCTCTCACGAGCCGCTTCTCAGGTCCGGCGGGCGTCTCCCTTGGGCTCACCCCCCGGGGCTGCCGTCGGCCAAAACCCCTG
TATCCAACCCCCTTTTTTAACACACCCCAAACCACAACCAACTCTGAAGCATCTTTGGTGGCCCGGCCCCGTGCCGTCCACTCCAAACCAAAGACAACTCTCAACA
ACGGATATCTTGGCTCCCGTATCGATGAGGAACGCAGCGAAATGCGATACGTAGTGTGAATTGCAGAATTCCGTGAACCATCGAATCTTTGAACGCAAATTGCG
CCCGAGGCTTCGGCCGAGGGCATGCCTGCCTCAGCGTCGGTTTACACCCTCGCCCTCCCCCACCGCTTGGCTGGGGTGCTGGTGCGGATCTGGCCCTCCCGGCT

CCGGCCCTGCCTTGTGCAGGGGCGCCCGGGTTGGCTGAAGCCCAGAGGCTTGAGCATGGACCCCGTTTGCAGGGCAATGGCTTGGTAGGTAGGCACCCCCTAC
GCAGCCTGCCGTTGCCCGAGGGGTCTTTGCTGGAGGCCCAGCAGGAATTCGGCCCTCACCGGCCGAACCACTCACTCATTCGACCT (GAGCTCAGGCAAGA)

#### >Ori_C46_C

(ACACACCGCCCGTCGCTCCTACCGATTGGGTGTGCTGGTGAAGTGTTCGGATTGGCAGCTTAGGGTGGCAACACCTCAGGTCTGCCGAGAAGTTCATTAAACC
CTCCCACCTAGAGGAAGGAGAAGTCGTAACAAGG)
TCTCCGTAGGTGAACCTGCGGAGGGATCATTGAATTATTAAAACCACAATGTGAACCTCAACGTTCCGTGCCCTGGCTTGCCAGTGGGGCGACATGGTCAACAC
CAGGTCGTACTCACAGCTGGGTGGGCATTGTTGCCTACTCAGTGGCGCCTTGGCATGATCATACACCAGTGCTAACCACTGATAAAACCAAACTCTGAAGTTTGA
TTGCTATTCATTGGCAATCTTAACCAAAGACAACTCTCAACAACGGATATCTTGGCTCCCGTATCGATGAAGAACGCAGCGAAATGCGATACGTAGTGTGAATTG
CAGAATTCCGTGAACCATCGAATCTTTGAACGCATATTGCGCTCGAGCCTTCGGGCAAGAGCATGTCTGCCTCAGCGTCGGTTTACACCCTCACCCCTCCCTTTCT
TGGGTGTGTTGATCTTTGATCAACCATTGGGGTGGATCTGGCTTCCCCAATCTGCCTTGTAGCGGATTGGGTTGGCTGAAGCACAGAGGCTTAAGCAAGGACCC
GATATGGGCTTCAACTGGATAGGTAGCAACGGCTTGTGCCGACTACACGAAGTTGTTGCCTGTGGACTTTGCTAGAGGCCAAGCAGGAACATGCTTATGCATGC
CTAAACTTTCGACCT (GAGCTCAGGCAAGG)

#### >SAG_211_8k_Chlorella_sorokiniana_owndata

(ACACACCGCCCGTCGCTCCTACCGATTGGGTGTGCTGGTGAAGTGTTCGGATTGGCGACCGGGGGCGGTCTCCGCTCTCGGCCGCCGAGAAGTTCATTAAACC
CTCCCACCTAGAGGAAGGAGAAGTCGTAACAAGG)
TTTCCGTAGGTGAACCTGCGGAAGGATCATTGAATCGATCGAATCCACACCGGTAACCACACTGTCGCCCTCGGCGGTGCATTCTCTGGCTTCGGCTGGGTTTCA
CCCCGAGCGTCGGCCCCTGGGTTGGGGTTCTCACGAGCCGCTCTCCAGGTCCGGCGGGCGCCTCCCTTGGGCTCACCCCCTGGGGCTGTCGTCGGCCAAAACCC
CTGTATCCAACCTTTTTTTTAACACACCCCAAACCACAACCAACTCTGAAGCATCTTTGGTGGCCCGGCCCCGTGCCGTCCACTCCAAACCAAAGACAACTCTCAA
CAACGGATATCTTGGCTCCCGTATCGATGAAGAACGCAGCGAAATGCGATACGTAGTGTGAATTGCAGAATTCCGTGAACCATCGAATCTTTGAACGCAAATTG
CGCCCGAGGCTTCGGCCGAGGGCATGTCTGCCTCAGCGTCGGTTTACACCCTCGCCCTCCCCCCCTGTGGGGGGCGGTGCGGACCTGGCCCTCCCGGCTCCGCT
CTCTCCCGAGCGTCCGGGTTGGCTGAAGCACAGAGGCTTGAGCATGGACCCCGTTTGTAGGGCAATGGCTTGGTAGGTAGGCACCCCCTACGCAGCCTGCCGT
TGCCCGAGGGGACTTTGCTGGAGGCCCAGCAGGAATCCGGCCCTTCCCGGCCGGACTACTCACTCATTCGACCT
(GAGCTCAGGCAAGA)

Comment: 100 % identity with UTEX1665; 1 base change with UTEX1230

#### >SAG211_34_owndata_GenBank_ MN194596

(ACACACCGCCCGTCGCTCCTACCGATTGGGTGTGCTGGTGAAGTGTTCGGATTGGCGACCGGGTGCGGTCTCCGCTCTCGGCCGCCGAGAAGTTCATTAAACC
CTCCCACCTAGAGGAAGGAGAAGTCGTAACAAGG)
TTTCCGTAGGTGAACCTGCGGAAGGATCATTGAATCGATCGAATCCACTCTGTGAACCAAACGTCCCCCCTTGGGTGCGGGCTTCGGTCTGCCCCAAGGCGTCG
GTTCCCTGGCTGGGGTCTTCGGACCGCAGTTAGGTCCGGCGGGCGCGCCCTCTGGCGTGTCGGCCCTCGTGGCTGCCGCCAGTTGGGTTCGCTGGAAATTGTAT
CCAACTCAACCCACCCCAAACCACAACTTATACTGAAGCAATCGGTGAGTGCACTCTGGTGCCTCGCTCTAACCAAAGACAACTCTCAACAACGGATATCTTGGC
TCCCGTATCGATGAAGAACGCAGCGAAATGCGATACGTAGTGTGAATTGCAGAATTCCGTGAACCATCGAATCTTTGAACGCAAATTGCGCCCAAGGCTTCGGC
CGAGGGCATGTCTGCCTCAGCGTCGGCTTACCCCCTCGCTCCCCCTCTCCTTTGGAGTGGGTGAACGGATCTGGTTTTCCCGGCTACGTGCTTCTGCACGCCCGG
GTTGACTGAAGTGTAGAGGCTTGAGCATGGACCCCGTTTGTAGGGCAATGGCTTGGTAGGTAGCTTAGCTACACCGCCTGCCGTGGTCCGAGGGGACTTTGCT
GGCGGCCCAGCAGGAATTCGGGTGTTGGGTTTCCCACCCCGAAAGCTTCAAACCTTCGACCT
(GAGCTCAGGCAAGA)

Comment: 100 % identity of sequence with data bank entries termed:
Chlorella sp. SAG 211-34 (clear)
Auxenochlorella pyrenoidosa (ambiguous)
Chlorella vulgaris (ambiguous)
Pseudochlorella pringsheimii (ambiguous)
Chlorella pyrenoidosa (ambiguous)

#### >AY591508_Chlorella_vulgaris_SAG_211-11b

TTTCCGTAGGTGAACCTGCGGAAGGATCATTGAATCTATCGAATCCACTTTGGTAACCACTCGTCCCCCTCGTCCGATGTCGCCCTCTCTTAGGAGA
GTGCGATGCGGCGAGCGTCGGTCCCCTGGCTGTGGCTCCCCCGAGCTGTTGCTCAGGTCCGGCGGGCGTCCCTTCACATGTGGGACCCCTTCTTTTT
GAGGGGACAATCCCCTTTGGAGGATCCGACGTCGGAAATTCCAACTCAACTCAACCCACCCCAAACTGAAACTTATTCTAAAGCACCTTGTGGTTGG
CAGCTCGTCTGCCGTCCACTCCAAACCAAATACAACTCTCAACAACGGATATCTTGGCTCCCGTATCGATGAAGAACGCAGCGAAATGCGATACGTA
GTGTGAATTGCAGAATTCCGTGAACCATCGAATCTTTGAACGCAACTTGCGCCTGAGGCTTCGGCCAAAGGCATGTCTGCCTCAGCGTCGGCTCACC
CCCTCGCCTCCCCATCTCATTTGATTGGGAAGGCGGATCTGACCTTCCCGGTTCCGCCGGTCACTCGTGATTGGCGCCGGGTCGGTTGAAGCTCAGA
GGTATGAGCATGGACCCCGTTCGTAGGGTAATGGCTTGGTAGGTAGGCATTCCCTACGCATCCTGCCGTTGCCCGAGGGGACTTTGCTGGAGACCTA
GCAGGAATTCGGATGCTTGGGCACCCCCCGACACCGAAACTCTTCATTTCGACCT
(GAGCTCAGGCAAGACTACCCGCTGAACTTAA)

#### Part IIb:

Sequence of sample C-126 (identical with 99.9 % identity with SAG 211-11b and UTEX 259

>C-126

```
(ACACACCGCCCGTCGCTCCTACCGATTGGGTGTGCTGGTGAAGTGTTCGGATTGGCGACCTGGGGCGGTCTCCGCTCTCGGCCGCCGAGAAGTTCA
TTAAACCCTCCCACCTAGAGGAAGGAGAAGTCGTAACAAGG)
TTTCCGTAGGTGAACCTGCGGAAGGATCATTGAATCTATCGAATCCACTTTGGTAACCACTCGTCCCCCTCGTCCGATGTCGCCCTCTCTTAGGAGA
GTGCGATGCGGCGAGCGTCGGTCCCCTGGCTGTGGCTCCCCCGAGCTGTTGCTCAGGTCCGGCGGGCGTCCCTTCACATGTGGGACCCCTTCTTTTT
GAGGGGACAATCCCCTTTGGAGGATCCGACGTCGGAAATTCCAACTCAACTCAACCCACCCCAAACTGAAACTTATTCTAAAGCACCTTGTGGTTGG
CAGCTCGTCTGCCGTCCACTCCAAACCAAATACAACTCTCAACAACGGATATCTTGGCTCCCGTATCGATGAAGAACGCAGCGAAATGCGATACGTA
GTGTGAATTGCAGAATTCCGTGAACCATCGAATCTTTGAACGCAACTTGCGCCTGAGGCTTCGGCCAAAGGCATGTCTGCCTCAGCGTCGGCTCACC
CCCTCGCCTCCCCATCTCATTTGATTGGGAAGGCGGATCTGACCTTCCCGGTTCCGCCGGTCACTCGTGATTGGCGCCGGGTCGGTTGAAGCTCAGA
GGTATGAGCATGGACCCCGTTCGTAGGGTAATGGCTTGGTAGGTAGGCATTCCCTACGCATCCTGCCGTTGCCCGAGGGGACTTTGCTGGAGACCTA
GCAGGAATTCGGATGCTTGGGCACCCCCCGACACCGAAACTCTTCATTCGACCT
(GAG)
```
