## Supplementary figures and images for "The wondrous and worrying diversity of the N‐glycans of *Chlorella* food supplements"

### 2a Figure S2 Chlorella products spectra grouped.pdf

#### Figure S2: Exemplary spectra of glyco-groups of Chlorella products. Order essentialy as in Table 1

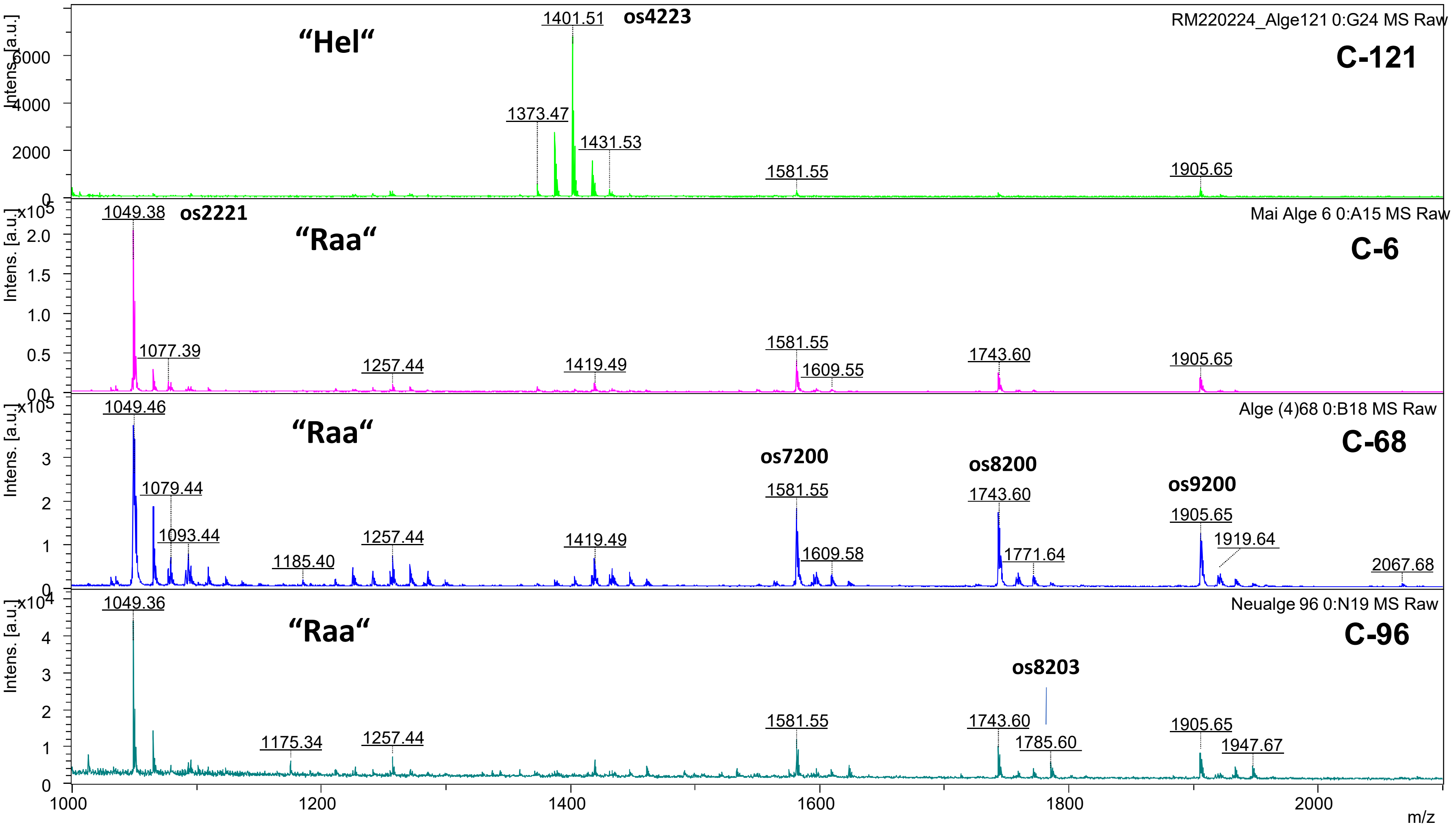

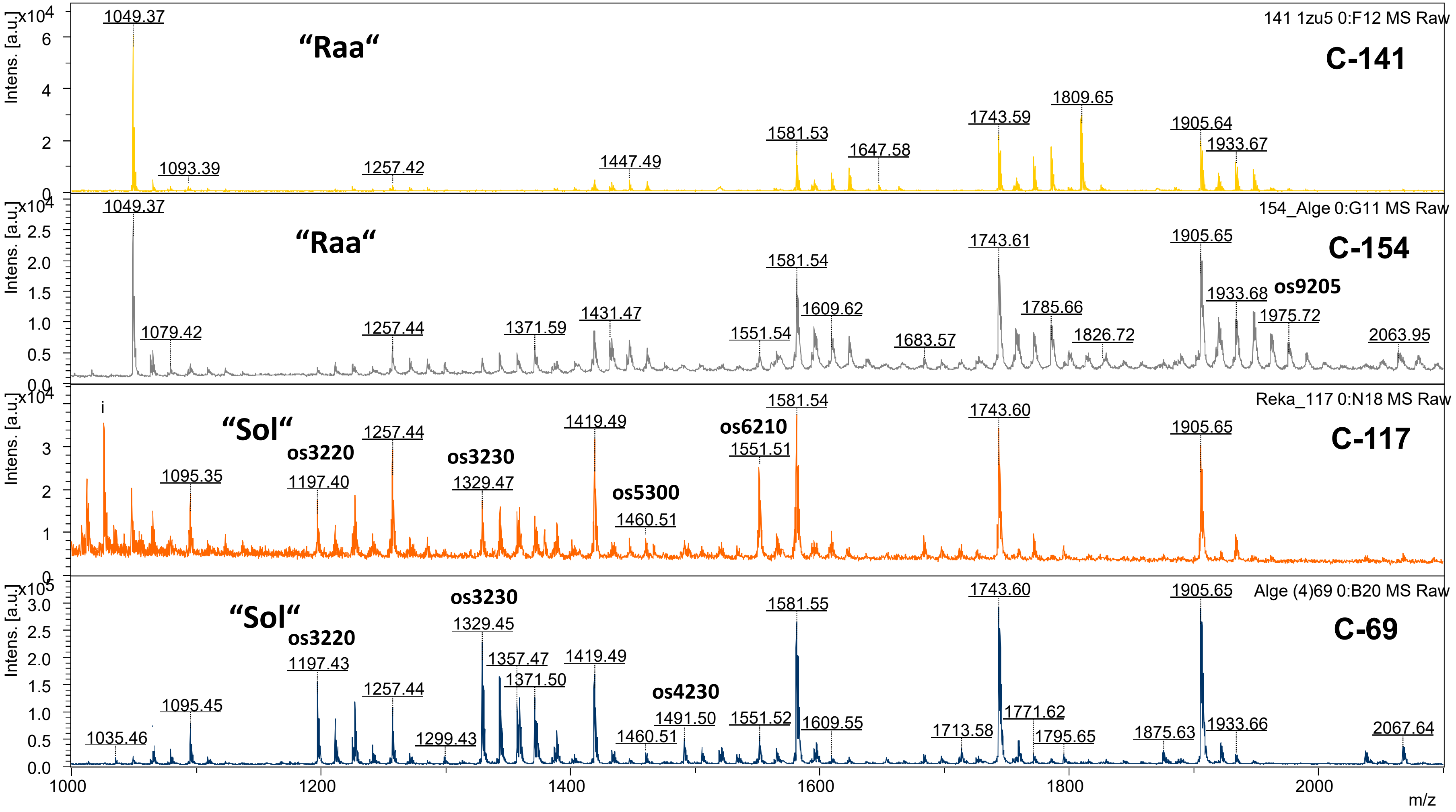

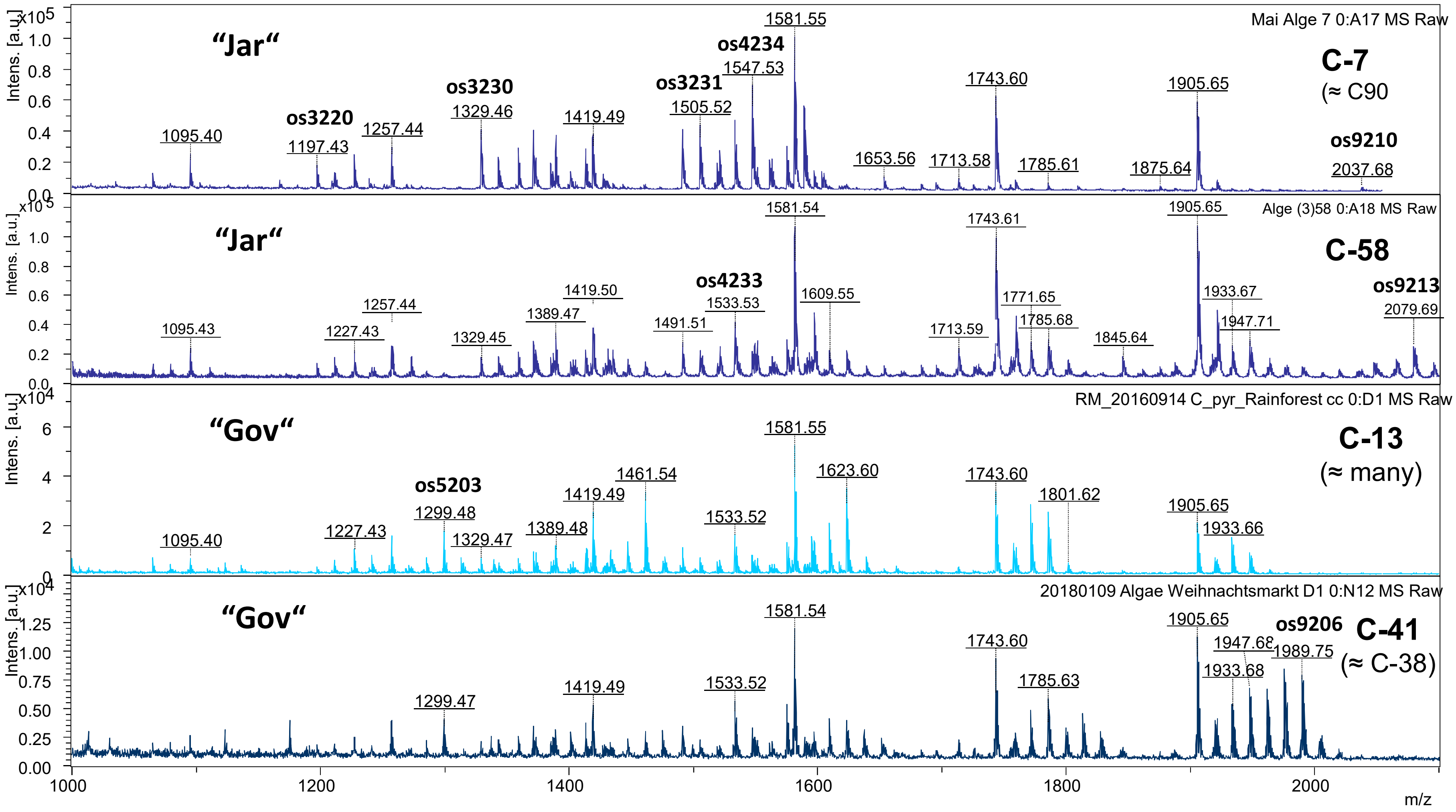

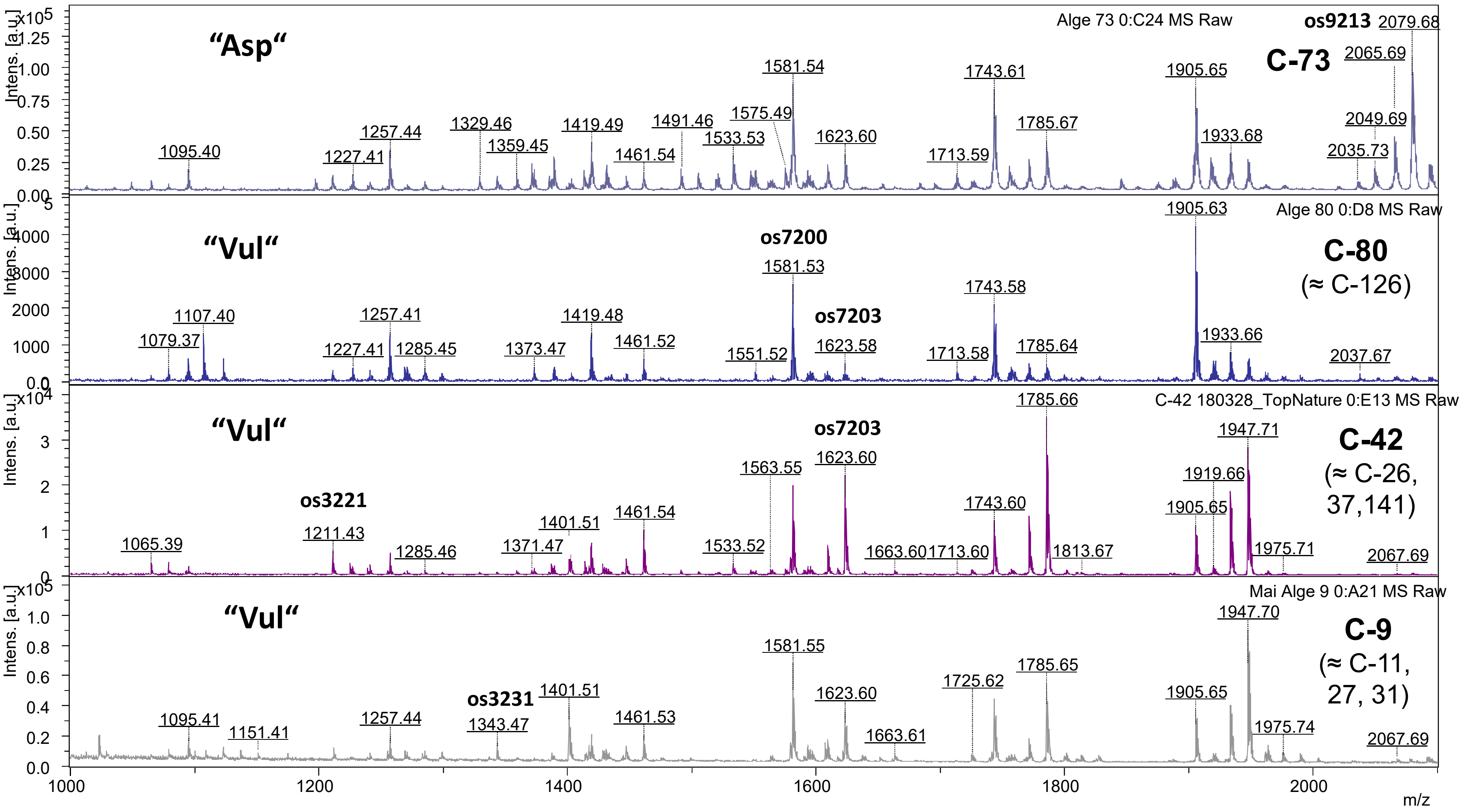

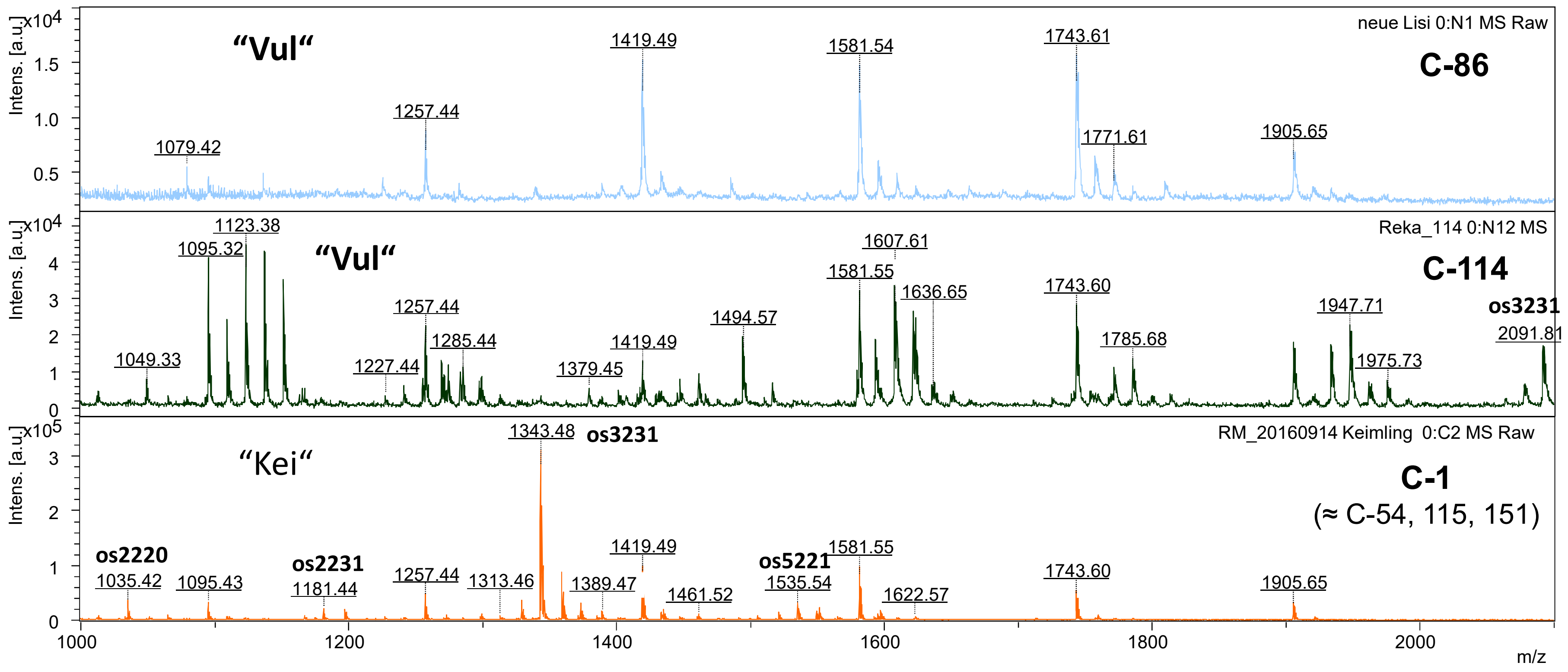
